# Inhibition of Soluble Epoxide Hydrolase Rescues Cognitive Deficits by Preserving Neurovascular Integrity and Attenuating Glial- and Neuropathology in Diabetic-Related Dementia

**DOI:** 10.64898/2026.06.01.729327

**Authors:** Xing Fang, Jane J. Border, Huawei Zhang, Gilbert C. Morgan, Andrew Gregory, Yvan Hanscom-Trofy, Rui Dong, Jun Yang, Sung Hee Hwang, Christophe Morisseau, Bruce D. Hammock, Fan Fan, Richard J. Roman

## Abstract

Diabetes mellitus (DM) is a major risk factor contributing to the development of Alzheimer’s disease-related dementias (ADRD). While one of the early symptoms of both Alzheimer’s disease (AD) and DM-related ADRD is a reduction in cerebral blood flow, the underlying biological mechanisms driving this decline remain to be fully elucidated. Genome-wide association studies have linked AD/ADRD to single-nucleotide polymorphisms in the gene encoding soluble epoxide hydrolase (sEH), an enzyme we previously reported to be upregulated in the brains of an AD rat model. Our previous work also demonstrated that chronic inhibition of sEH with 1-trifluoromethoxyphenyl-3-(1-propionylpiperidin-4-yl) urea (TPPU) preserves hippocampal-dependent spatial learning and memory and improves cerebral hemodynamics in both AD and DM-ADRD models. In the present study, we found that chronic TPPU treatment (1 mg/kg/day for 9 weeks) reduced brain sEH expression, improved cortical-based long-term non-spatial recognition memory involving both cortical and hippocampal networks, and reduced anxiety in DM-ADRD rats. TPPU improved brain perfusion and normalized impaired whisker-evoked functional hyperemia, an effect linked to upregulation of Kir2.1 expression in cerebral capillaries. Furthermore, TPPU restored tight junction proteins (ZO-1 and OCLN), mitigated capillary rarefaction, and suppressed astrocyte and microglial activation. At the cellular level, TPPU attenuated hippocampal neurodegeneration, restored the expression of synaptic proteins (PSD95 and SY38), and reduced levels of key pro-inflammatory chemokines, including MCP-1, RANTES, and MIP-1α, in DM-ADRD. In conclusion, TPPU preserves cognitive function in DM-ADRD by mitigating cerebrovascular dysfunction, neuroinflammation, and gliosis while protecting synaptic integrity and neuronal survival, representing a promising therapeutic strategy for DM-ADRD.

## INTRODUCTION

Diabetes mellitus (DM) is a major risk factor contributing to the development of Alzheimer’s disease-related dementias (ADRD).^1-3^ While one of the early symptoms of both Alzheimer’s disease (AD) and DM-related ADRD is a reduction in cerebral blood flow (CBF), the underlying biological mechanisms driving this decline remain to be fully elucidated, despite evidence that impaired brain perfusion precedes the onset of AD pathologies and cognitive deficits.^4-7^

Cerebral hypoperfusion in AD/ADRD is driven by a multifaceted dysfunction of the vascular and glymphatic systems in the brain.^8-10^ At the foundational level, CBF autoregulation is compromised when vascular smooth muscle cells (VSMCs) or contractile pericytes fail to mount an appropriate myogenic response, leaving the brain vulnerable to fluctuations in systemic blood pressure.^11-14^ This is compounded by the loss of neurovascular coupling, a mechanism that normally facilitates functional hyperemia, which increases blood flow to active neurons. When this process is impaired, high-demand brain regions become oxygen-deprived.^15,16^ Crucial to this environment is the blood-brain barrier (BBB), in which structural decay of the endothelium, loss of pericytes, and breakdown of tight junctions (TJs) lead to microvascular leakage and neurotoxicity. ^6,17,18^ Within the microvasculature, capillary stalling occurs when endothelial dysfunction triggers chemokine release and adhesion molecule expression, leading to leukocyte plugging of small vessels and further reducing perfusion.^19,20^ Additionally, the waste-removal machinery of the brain fails when glymphatic drainage is disrupted; this is often marked by astrocyte activation, depolarization of astrocytic endfeet, and mislocalization of aquaporin-4 (AQP4) water channels, thereby preventing the efficient clearance of toxic metabolites such as amyloid beta (Aβ).^10,21,22^

The decline of the vascular and glymphatic systems in AD/ADRD is driven by synergistic biochemical and genetic insults that progressively impair cerebral homeostasis. Chronic neuroinflammation and oxidative stress accelerate endothelial and pericyte dysfunction, weaken BBB integrity, and promote immune-mediated capillary obstruction.^14^ Genetic risk factors, including the APOE ε4 allele and variants in genes encoding soluble epoxide hydrolase (sEH), ^23-25^ COX-2,^26^ CYP4A11, CYP4F2,^11,27-29^ and ADD3,^30,31^ have also been shown to increase neurovascular vulnerability. As vascular and glymphatic function deteriorates, impaired clearance of Aβ and other proteotoxic metabolites drives a self-reinforcing cycle in which toxic aggregates accumulate within perivascular spaces and vessel walls, worsening neurovascular uncoupling and cerebral hypoperfusion.

Current therapeutic development for AD/ADRD is increasingly shifting beyond neuron-centric models toward integrated strategies targeting neurovascular dysfunction, glymphatic impairment, and chronic neuroinflammation.^32^ Anti-inflammatory biologics, such as ANX005 (tanruprubart), a C1q complement inhibitor, are being explored to suppress microglial-driven vascular inflammation and synaptic injury,^33^ while antioxidant and vasoprotective agents, including edaravone,^34^ N-acetylcysteine,^35^ sEH inhibitors,^24,36-38^ and sEH/COX-2 dual inhibitors,^39^ aim to reduce endothelial dysfunction, oxidative stress, and pericyte degeneration within the neurovascular unit. Therapeutic modulation of AQP4 localization and glymphatic transport has also emerged as a promising strategy for treating neurodegenerative diseases. While the critical role of this pathway is frequently validated using TGN-020, an AQP4 inhibitor that blocks glymphatic flow and worsens protein aggregation,^40^ AQP4 facilitators such as TGN-073 are currently being investigated for their potential to enhance Aβ and tau clearance.^41^ Although many of these therapies remain in preclinical or early clinical stages, they collectively reflect a major paradigm shift toward restoring vascular integrity, immune homeostasis, and brain clearance systems as disease-modifying approaches for AD/ADRD.

We previously reported that sEH protein expression is upregulated in the brain in the AD rat model.^38^ Inhibition of sEH with 1-trifluoromethoxyphenyl-3-(1-propionylpiperidin-4-yl) urea (TPPU) improved cerebral hemodynamics and hippocampal-based spatial learning and memory in both AD and DM-ADRD rat models.^24,38^ Chronic TPPU treatment did not reduce body weight in either AD or DM-ADRD rats; however, it significantly reduced plasma glucose and HbA1c levels only in the DM-ADRD model, although glycemia remained within the diabetic range and was not normalized.^24^ Transcriptomic profiles of primary cerebral VSMCs and hippocampal tissues demonstrated that TPPU treatment in AD rats reduced inflammation and oxidative stress, and was associated with reduced Aβ plaque burden, BBB leakage, glial activation, and neurodegeneration.^38,42^ Moreover, dual inhibition of sEH and COX-2 had a similar effect in improving cerebral vascular and cognitive function in AD rats, reducing vascular inflammation, and attenuating vascular remodeling.^39^ In related studies, we have reported that inhibition of sodium-glucose cotransporter-2 (SGLT2) with luseogliflozin attenuated mitochondrial dysfunction and oxidative stress in cerebral VSMCs, restored CBF autoregulation and functional hyperemia, reduced BBB leakage and neurodegeneration, and improved cognitive performance in both AD^43^ and DM-ADRD rat models.^44^ Notably, metabolic effects were observed only in the DM-ADRD model, where SGLT2 inhibition reduced body weight, plasma glucose, and HbA1c, effectively lowering glucose toward the normoglycemic range, whereas no significant metabolic changes were detected in the non-diabetic AD model.

The involvement of sEH in diabetes and cognitive decline with aging has been increasingly recognized.^24,45,46^ Epoxyeicosatrienoic acids (EETs) and other epoxy fatty acids (EpFAs) derived from the metabolism of fatty acids via CYP450 2C pathways possess potent anti-inflammatory properties in addition to their vasodilatory effects.^47^ However, EETs are rapidly metabolized by sEH to less active dihydroxyeicosatrienoic acids (DHETs).^48^ sEH is widely expressed in neurons, vascular ECs, SMCs, astrocytes, microglia, oligodendrocytes, adventitial cells, and ependymal cells in the central nervous system, ^49,50^ and the expression of sEH is elevated in patients and rodent models of diabetes.^46,51^ These findings suggest that inhibiting sEH or administering EpFA agonists may delay the development of age-related cognitive dysfunction in diabetes.^45,46,52,53^ Building upon our previous findings that TPPU improved hippocampal-dependent spatial learning and memory using the eight-arm water maze in DM-ADRD, ^24^ we expanded the behavioral assessment in the present study to include long-term non-spatial recognition memory involving both cortical and hippocampal networks, as well as anxiety-like behaviors using Novel Object Recognition (NOR) and Open Field Tests. Complementing these cognitive assessments, we extended our *ex vivo* pressure myography investigations, following up on our previous discovery that the myogenic responses of middle cerebral arteries (MCAs) and parenchymal arterioles (PAs) were restored in TPPU-treated DM-ADRD rats.^24^ We also evaluated myogenic tone and absolute inner diameters (IDs) across a wide spectrum of perfusion pressures below and above the autoregulatory range. Finally, to link the structural microvascular properties to functional blood flow, we evaluated baseline cortical perfusion and whisker-evoked functional hyperemia. Furthermore, we sought to elucidate the underlying molecular mechanisms, focusing on capillary Kir2.1 expression, tight junction protein integrity, and synaptic markers. Finally, we assessed whether sEH inhibition could mitigate the chronic neuroinflammatory environment by quantifying glial activation and the expression of pro-inflammatory cytokines and chemokines.

## MATERIALS AND METHODS

### General

Experiments were performed in male T2DN rats and non-diabetic control (Ctrl) Sprague-Dawley (SD) rats. We utilized the T2DN strain as a model for DM-ADRD, as our previous reports have established that these rats develop chronic type 2 diabetes along with significant cerebrovascular and cognitive dysfunction.^6,14,17,44^ The rats were bred and housed at the University of Mississippi Medical Center (UMMC) and Augusta University and had free access to water and diet. TPPU was initially dissolved in 100% polyethylene glycol 400 (PEG-400; 91893, MilliporeSigma, Burlington, MA) and subsequently diluted in drinking water to a final PEG-400 concentration of 1%. TPPU was synthesized by Dr. Hammock’s team.^54^ The T2DN rats received either a vehicle (1% PEG-400 in drinking water) or TPPU (1 mg/kg/day in the vehicle) starting at 15 months of age for 9 weeks. ^24,38,55^ Experimental design and sequential cohort allocation are presented in **Figure 1**. The present study was conducted in accordance with the National Institutes of Health Guide for the Care and Use of Laboratory Animals. All protocols were approved by the Institutional Animal Care and Use Committee of UMMC and Augusta University.

**Figure 1.**
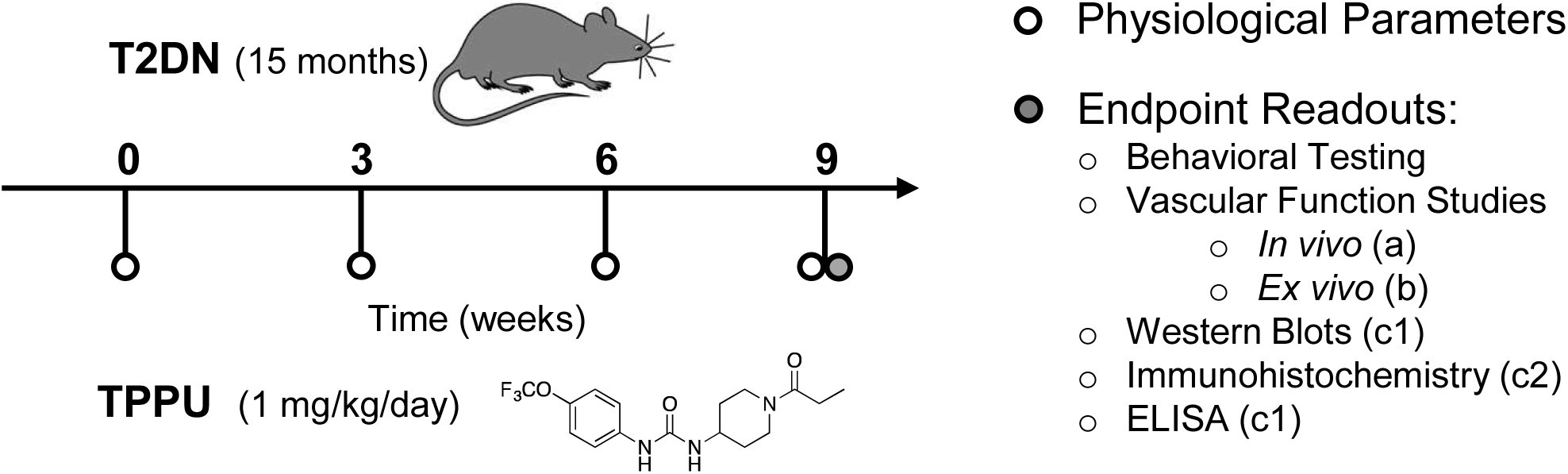
Experimental Design and Sequential Cohort Allocation. Schematic timeline of the 9-week study protocol. T2DN and age-matched control rats received either vehicle (1% PEG-400 in drinking water) or the soluble epoxide hydrolase inhibitor TPPU (1 mg/kg/day in vehicle) for 9 weeks, starting at 15 months of age. Physiological parameters (body weight, blood glucose, and HbA1c) were monitored every 3 weeks. To accommodate the extensive terminal assays, the study was conducted in three independent, sequential cohorts maintained under identical experimental conditions. A battery of behavioral tests, including Novel Object Recognition, Open Field, and the Eight-arm Water Maze, were performed at the 9-week endpoint. Following behavioral assessment, rats were processed for specific endpoint readouts: (a) *in vivo* vascular function; (b) *ex vivo* vascular function; and (c1) protein expression (Western blot) and ELISA, (c2) neuropathological assessment (Immunohistochemistry).

### Behavioral Tests

#### Novel Object Recognition Test

The NOR test was used to evaluate long-term non-spatial recognition memory, a process requiring consolidated cortical and hippocampal memory. The protocol was executed across three distinct phases: habituation, familiarization, and testing. During the habituation phase, rats were placed individually into the empty open field arena and allowed to freely explore for 10 minutes. Twenty-four hours (24 h) later, the familiarization phase was conducted. Two identical familiar objects were placed in opposite corners of the arena, and each rat was allowed to explore them for 10 minutes. Following a 4-hour retention delay in their home cages, rats underwent the test phase. During this 10-minute session, one of the familiar objects was replaced with a physically distinct novel object. To prevent confounding spatial or object preference biases, both the location of the novel object (left versus right corner) and the specific objects designated as familiar or novel stimuli were counterbalanced across groups. The duration of time spent actively exploring each object was recorded.

Memory performance was quantified using the following indices:

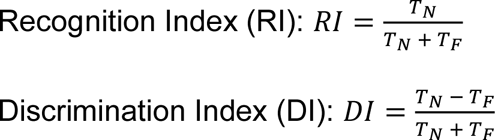

Where *TN* represents the time spent exploring the novel object during the testing phase, and *TF* represents the time spent exploring the familiar object. Positive DI values indicate a preference for the novel object, negative values indicate a preference for the familiar object, and a value of zero indicates no preference. A failure to preferentially explore the novel object was interpreted as a long-term recognition memory deficit. Between all trials and animals, the apparatus and objects were thoroughly sanitized with 70% ethanol and rinsed with distilled water to eliminate confounding olfactory cues.

#### Open Field Test

The open field test was used to assess locomotor activity, exploratory behavior, and anxiety-like behavior, as previously described.^6^ Rats were acclimated to the testing room for 2 hours prior to the start of the procedure. Each animal was placed in the center of a clear plastic chamber (MED-OFAS-RSU, Med Associates, St. Albans, VT) equipped with the Opto-Varimex ATM3 Auto-Track System (Columbus Instruments, Columbus, OH). This system utilized an array of infrared photobeams to monitor movement via beam breaks. Activity was recorded over a 30-minute period to quantify total distance traveled and resting time as measures of general motor function. To assess anxiety-like behavior, the system tracked the spatial distribution of the rat, specifically comparing time spent in the center of the arena versus the periphery (thigmotaxis).

### *Ex Vivo* Cerebrovascular Pressure Myography

#### Vessel Isolation and Cannulation

The MCAs and PAs were isolated based on our established protocols.^11,42,56,57^ Briefly, rats were euthanized under 4% isoflurane anesthesia. Brains were rapidly harvested into ice-cold, calcium-free physiological salt solution (PSS_0Ca_) containing 119 NaCl, 4.7 KCl, 1.17 MgSO_4_, 0.03 EDTA, 18 NaHCO3, 5 HEPES, 1.18 NaH_2_PO_4_, and 10 glucose (all in mM; pH 7.4). A cortical section was dissected, and branch-free M2 segments of the MCAs and downstream PAs were then dissected in ice-cold PSS_0Ca_ supplemented with 1% bovine serum albumin. Vessel segments were secured onto glass micropipettes (1B120-6, World Precision Instruments) in a pressure myograph chamber (Living Systems Instrumentation, Burlington, VT). The chamber was maintained at 37°C and continuously perfused with aerated (21% O_2_, 5% CO_2_, and 74% N_2_) PSS_Ca_ to maintain pH 7.4. PSS_Ca_ was prepared with the same composition as PSS0Ca, but without EDTA and with 1.6 mM CaCl_2_. Following cannulation, vessels were maintained at their *in situ* length, pressurized to 40 mmHg for MCAs and 10 mmHg for PAs, and equilibrated for 30 minutes to develop spontaneous tone. Vascular viability was confirmed by a >15% vasoconstrictive response to 60 mM KCl; vessels failing this criterion were excluded.

#### Absolute Inner Diameter Measurements

After preconditioning, IDs were measured across a wide range of perfusion pressures to assess the structural and active responses of the cerebral vasculature. IDs were recorded over a range from 40 to 180 mmHg in 20-mmHg increments and from 10 to 60 mmHg in 10-mmHg increments for the MCA and the PA, respectively. After active measurements, the vessels were thoroughly rinsed with PSS_0Ca_ to determine the passive inner diameter (ID_0Ca_) at each corresponding pressure point. Measurements were obtained for both active in PSS_Ca_ (as ID_Ca_) and passive in PSS_0Ca_ conditions.

#### Myogenic Tone Quantification

To quantify the capacity of the vessels to constrict in response to increased intraluminal pressure, myogenic tone was calculated for both the MCA and the PA. Myogenic tone was expressed as a percentage of the passive diameter using the following formula:

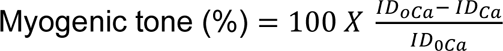

### *Ex Vivo* Functional Pressurized Capillary-Parenchymal Arteriole (CaPA) Preparation

#### CaPA Isolation and Cannulation

Following euthanasia with isoflurane and rapid decapitation, the brain was quickly harvested and immersed in ice-cold PSS_0Ca._ Under a stereomicroscope, a CaPA network was carefully microdissected from the parenchymal tissue using ultra-sharp forceps to isolate a feed parenchymal arteriole along with its downstream capillary branches. The isolated microvascular network was transferred to a pressure myography chamber (Living System Instrumentation, Burlington, VT), where the proximal end of the feeding arteriole was carefully cannulated onto a glass micropipette and secured with microsutures. To achieve a closed, pressurized system, open side branches were occluded with secondary sutures, and the distal ends of the capillary branches were self-sealed or gently compressed against the glass coverslip. Vessels were maintained at their *in situ* length, pressurized to 10 mmHg, and equilibrated for 30 minutes to develop spontaneous tone. Vascular viability was confirmed by a vasoconstrictive response >15% to 60 mM KCl; vessels failing this criterion were excluded.

#### Potassium-Induced Dilation

To mimic physiological neurovascular coupling responses, PSS_Ca_ containing 10 mM KCl was applied to the distal capillary end using a micromanipulator and glass micropipette. The resulting retrograde hyperpolarization signal was continuously tracked as percentage changes in IDs of the upstream feed PA over a 2-minute period.

#### Kir2.1 Channel Blockade

To evaluate inward-rectifier potassium channel dependency, PSS_Ca_ containing the selective Kir2.1 blocker ML133 (10 μM) was subsequently micro-injected near the terminus of the capillary following the assessment of the initial potassium-induced response. The subsequent changes in the upstream PA IDs were recorded over a 5-minute period.

### *In Vivo* Laser Speckle Contrast Imaging (LSCI)

#### Surgical Preparation

Rats were anesthetized using a combination of inactin (100 mg/kg, *i.p*.) and ketamine (60 mg/kg, *i.m*.). Body temperature was maintained at 37 °C. A small thinned cranial window was prepared using a low-speed dental drill under continuous saline cooling.

#### Resting Brain Perfusion Assessment

Spatially resolved regional cerebral blood flow and resting baseline brain perfusion maps were acquired through the intact, thinned bone window using an LSCI system. Resting perfusion values were calculated over a stable baseline phase and expressed as a percentage of the average perfusion level measured relative to the controls.

#### Whisker Stimulation-Induced Functional Hyperemia

To evaluate neurovascular coupling kinetics, mechanical stimulation was applied to the contralateral vibrissae by synchronously deflecting the whiskers at 10 Hz for 60 seconds. Cortical microvascular perfusion changes within the somatosensory domain were tracked continuously throughout a pre-stimulation baseline phase, the active stimulation window, and a post-stimulation recovery phase. Functional hyperemia responses were quantified as percentage changes from the pre-stimulation baseline.

#### Cerebral Autoregulation and Vascular Integrity

To assess the stability of cerebral perfusion across varying systemic pressures, mean arterial pressure (MAP) was pharmacologically manipulated. MAP was elevated incrementally with graded intravenous infusions of phenylephrine or decreased by controlled blood withdrawal as we previously reported.^6,44^ LSCI was used to generate real-time spatiotemporal maps of cortical surface perfusion at pressures of 60, 100, and 160 mmHg, representing values below, within, and above autoregulatory breakthrough points we previously identified in this DM-ADRD model.^6,44^ Vascular integrity and potential BBB leakage were qualitatively assessed by analyzing the extravasation of fluorescent tracers or the emergence of localized "hot spots" of high perfusion in LSCI maps at hypertensive pressures.

### Western Blot Analysis

As previously described, one brain hemisphere from each rat was homogenized in ice-cold RIPA lysis buffer supplemented with a 1% protease and phosphatase inhibitor cocktail (A32963; Thermo Fisher Scientific, Waltham, MA). Initial homogenization was performed using a ground-glass homogenizer, followed by mechanical disruption with a FastPrep-24 bead homogenizer (15070599; MP Biomedicals, Santa Ana, CA). The homogenate was centrifuged at 14,000 × g for 15 min at 4°C to pellet insoluble cellular debris. Total protein concentration in the supernatant was determined via a Bradford assay (5000006; Bio-Rad Laboratories, Hercules, CA) using bovine serum albumin (BSA) as the standard. Equal amounts of protein were resolved on 4 - 20% Criterion™ TGX Stain-Free™ precast gels (5678093; Bio-Rad). Total protein loading per lane was visualized and captured under UV illumination using a ChemiDoc XRS+ imaging system (1708265; Bio-Rad) to serve as a normalization control. Proteins were subsequently transferred to nitrocellulose membranes (1704159; Bio-Rad) using a Trans-Blot Turbo Transfer System (690BR008061; Bio-Rad). Membranes were blocked in Tris-buffered saline containing 0.1% Tween-20 (TBST) and 5% non-fat dry milk for 2 hours at room temperature, followed by incubation with primary antibodies overnight at 4°C. Following four washes with TBST, membranes were incubated with horseradish peroxidase (HRP)-conjugated secondary antibodies for 2 hours at room temperature, followed by final washes in TBST. Protein bands were detected using SuperSignal™ West Dura Extended Duration Substrate (34075; Thermo Fisher Scientific), imaged on the ChemiDoc XRS+ system, and quantified using densitometry normalized against the total lane protein signal.

The primary antibodies utilized were: mouse anti-sEH (sc-166961, 1:1000; Santa Cruz Biotechnology, Dallas, TX), rabbit anti-ZO-1 (617300, 1:1000; Thermo Fisher Scientific), rabbit anti-occludin (SAB5700784, 1:1000; MilliporeSigma, Burlington, MA), rabbit anti-GFAP (AB5804, 1:1000; MilliporeSigma), rabbit anti-IBA1 (016-20001, 1:1000; FUJIFILM Wako Pure Chemical, Osaka, Japan), mouse anti-synaptophysin [SY38] (ab8049, 1:1000; Abcam, Cambridge, UK), and mouse anti-PSD95 (MAB1598, 1:1000; MilliporeSigma). The secondary antibodies included HRP-conjugated goat anti-rabbit IgG (ab6721, 1:5000; Abcam) and HRP-conjugated rabbit anti-mouse IgG (ab97046, 1:5000; Abcam).

### Immunohistochemistry

As previously described, freshly collected brain hemispheres were fixed in a 10% zinc-formalin solution for 48 hours and subsequently stored in phosphate-buffered saline (PBS) containing 1% sodium azide at 4°C until processing. Free-floating coronal tissue sections (30 µm) were prepared using a Compresstome vibrating microtome (VF-310-0Z; Precisionary Instruments, Ashland, MA) following embedding in low-melting agarose. Sections were washed three times for 5 minutes each, incubated in Dulbecco’s PBS containing 5% BSA and 2% Triton X-100 for 1 hour at room temperature, and then incubated overnight at 4°C with primary antibodies against rabbit glial fibrillary acidic protein (GFAP; AB5804, 1:1000; MilliporeSigma) or (on separate sections) rabbit ionized calcium-binding adapter molecule 1 (Iba1; 019-19741, 1:500; FUJIFILM Wako). The following day, sections were washed and incubated with an Alexa Fluor 555-conjugated goat anti-rabbit IgG secondary antibody (A21428, 1:1000; Invitrogen) for 2 hours at room temperature. Sections were then counterstained with NeuroTrace 500/525 Green fluorescent Nissl stain (N21480, 1:200; Thermo Fisher Scientific) for 2 hours at room temperature. After the final washes, the sections were mounted on glass slides and coverslipped using ClearVue mounting medium (4212; Epredia, Portsmouth, NH). For separate staining protocols, sections were incubated with NeuroTrace 530/615 Red fluorescent Nissl stain (N21482, 1:200; Thermo Fisher Scientific) diluted directly in a blocking solution (5% BSA and 2% Triton X-100 in DPBS) for 2 hours at room temperature, followed by incubation with fluorescein-labeled tomato lectin (L32470, 1:200; Thermo Fisher Scientific) for 48 hours at 4°C.

To evaluate the microvascular coverage of Kir2.1 channels as previously described, separate rat brains were collected, fixed in 4% paraformaldehyde overnight at 4°C, and consecutively cryoprotected in 10% and 30% sucrose solutions at 4°C overnight. One hemisphere was embedded in Tissue-Tek® O.C.T. compound (4583; Sakura Finetek USA, Torrance, CA) and frozen. Coronal sections (50 µm) were prepared from the frozen tissue blocks using a CryoStar™ NX50 cryostat (Thermo Fisher Scientific). Free-floating frozen sections were collected and subjected to antigen retrieval by incubation with a 0.1% trypsin solution at 37°C for 15 minutes, followed by heating in a 10 mM sodium citrate buffer (pH 6.0) at 80°C for 3.5 hours. Following three washes, sections were incubated for 72 hours at 4°C with primary antibodies against mouse anti-collagen IV (Col IV; 1:200, MA1-22148; Thermo Fisher Scientific) and rabbit anti-Kir2.1 (1:200, APC-159; Alomone Labs, Jerusalem, Israel) diluted in the same blocking solution mentioned earlier. Sections were washed three times and subsequently incubated overnight at 4°C with Alexa Fluor 488–conjugated anti-mouse and Alexa Fluor 555–conjugated anti-rabbit secondary antibodies (Thermo Fisher Scientific) diluted in the same blocking solution. Following final washes, the sections were mounted on glass slides, treated with a VECTASHIELD antifading mounting medium with DAPI (H-1500-10, Vector Laboratories, Burlingame, CA), and finally coverslipped.

Confocal images for Kir2.1 coverage were acquired using a Nikon C2+ laser scanning confocal microscope on an Eclipse Ti2 inverted platform. Epifluorescence images for glial activation, neurodegeneration, and capillary density quantification were captured using a Nikon Eclipse 55i upright microscope equipped with a Nikon DS-Df1 digital color camera and NIS-Elements imaging software (v4.6). Glial reactivity was determined by counting the number of morphologically activated astrocytes and microglia per field across 2 sections and 2–3 distinct regions of interest within the hippocampal CA1 and CA3 subfields (7-8 rats per group). Capillary density was quantified as the area fraction of lectin-positive microvessels across 2–3 fields per hippocampal subfield (8-9 rats per group). Image processing and quantitative morphometric analyses were performed using ImageJ software (NIH, Bethesda, MD).

### Assessment of Brain Inflammatory Markers

Brain tissue (∼100 mg) was homogenized in cell lysis buffer (EA-0001, Signosis, USA) using a FastPrep-24 system with Lysing Matrix B tubes, followed by brief sonication on ice. Homogenates were centrifuged at 12,000 × g for 15 minutes at 4 °C, and supernatants were collected. Total protein concentration was quantified via a Bradford assay. Samples (20 µg/100 µL per well) were loaded onto a Rat Inflammatory Cytokine ELISA Plate Array strip (ES-1201, Signosis) pre-coated with antibodies against targeted cytokines and chemokines. Following a 2-hour incubation at room temperature, plates were washed three times and incubated with a biotin-labeled secondary antibody for 1 hour. After three additional washes, a streptavidin-HRP solution was applied for 45 minutes. The wells were washed three times, incubated with a chromogenic substrate for 30 minutes in the dark, and neutralized with the stop solution. Absorbance was recorded at 450 nm using a microplate reader, and values were expressed as fold change relative to controls.

### Statistical Analysis

All data are presented as mean values ± standard error of the mean (SEM). Statistical analyses were performed using GraphPad Prism 10 (GraphPad Software, Inc., Boston, MA, USA). For single-point comparisons involving a single independent variable across three groups, significance was determined using a one-way analysis of variance (ANOVA) followed by Tukey’s post-hoc test. For comparisons involving multiple factors or independent parameters across groups (such as distinct brain regions, proteins, or time-course assessments), data were analyzed using a two-way ANOVA or two-way repeated-measures ANOVA, followed by either Sidak’s post hoc multiple comparisons tests, as appropriate. A *p*-value < 0.05 was considered statistically significant.

## RESULTS

We first confirmed our previous findings^24^ that chronic TPPU treatment improved non-spatial short-term memory and spatial learning and memory in the eight-arm water maze while significantly reducing plasma glucose and HbA1c levels in the DM-ADRD model, although glycemia remained within the diabetic range and body weight was unaffected (confirmation data not shown).

### Chronic TPPU Treatment Attenuates Upregulation of sEH Protein in the Brain of DM-ADRD Rats

To investigate sEH expression in the brain, Western blot analysis was performed on total brain tissue homogenates (**Figure 2**; n = 9 per group). The sEH expression was significantly higher in the brains of elderly DM rats compared to age-matched SD controls (1.63 ± 0.08- vs. 1.00 ± 0.08-fold). Chronic TPPU treatment significantly reduced brain sEH expression to 1.14 ± 0.03-fold, effectively returning sEH protein levels to a range statistically indistinguishable from the control group. These findings demonstrate that TPPU treatment effectively prevents the increase in sEH protein levels in the brain of DM-ADRD rats.

**Figure 2.**
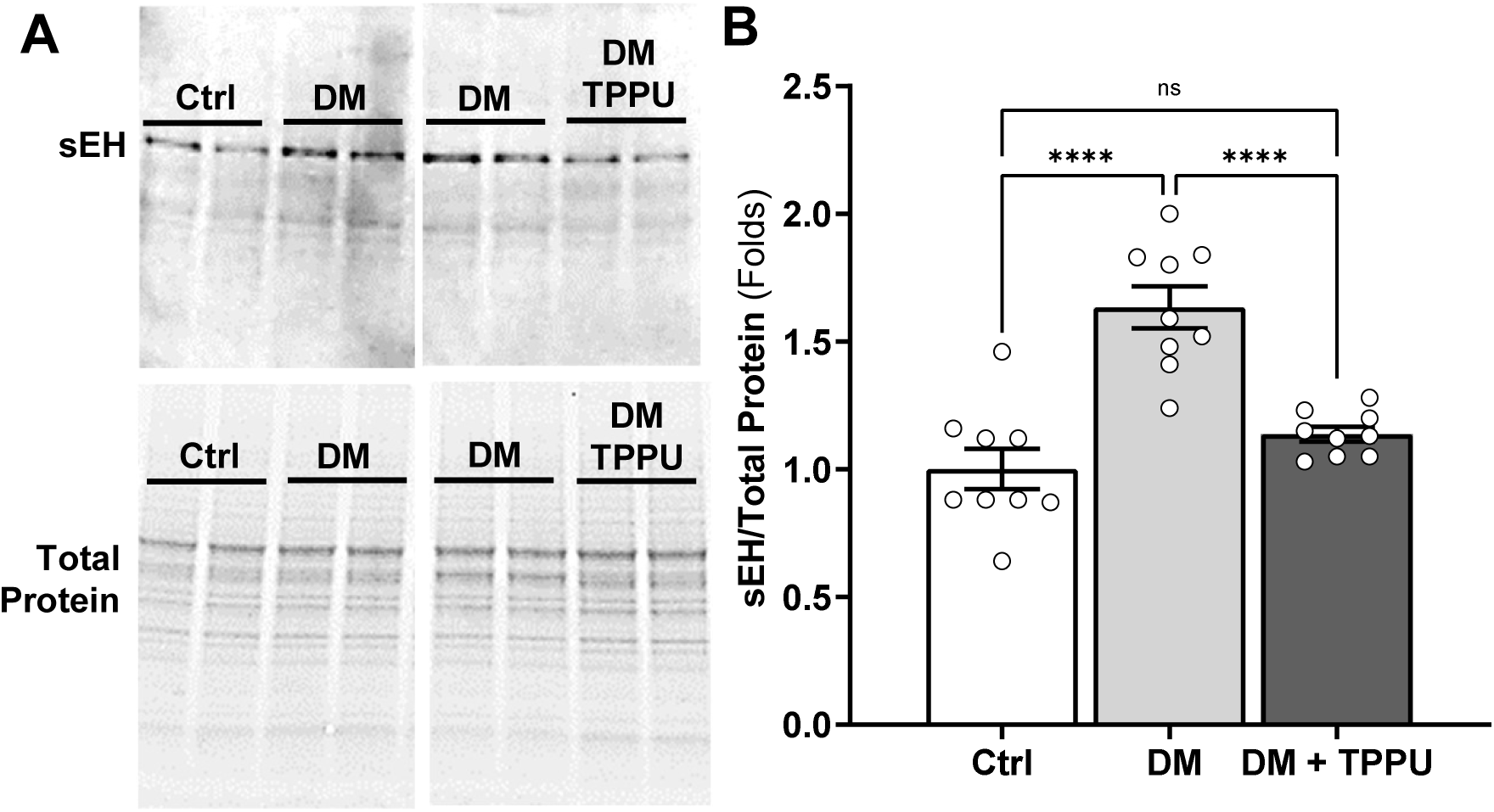
TPPU Treatment Attenuates Diabetes-Induced Upregulation of sEH in Total Brain Tissue. Representative Western blot images showing sEH protein expression in total brain tissue homogenates from **(A)** DM vs. Control (Ctrl) and DM + TPPU vs. DM. The top bands represent sEH (62 kD), while the bottom panels display the corresponding total protein loading control used for normalization. **(B)** Densitometric quantification of sEH protein levels normalized to total protein. Data are expressed as mean ± SEM (n = 9 rats per group). Significant differences were determined by one-way ANOVA followed by Tukey’s post-hoc test. \*\*\*\**p* < 0.0001; ns, non-significant.

### Chronic TPPU Treatment Improves Cognitive Performance in DM-ADRD Rats

NOR (**Figures 3A-B**) was performed to evaluate the impact of chronic sEH inhibition on long-term non-spatial recognition memory. Untreated DM-ADRD rats exhibited significant memory impairment compared with age-matched controls, as evidenced by a lower recognition index (0.34 ± 0.04 vs. 0.61 ± 0.05) and a negative discrimination index (−0.32 ± 0.08 vs. 0.21 ± 0.09). Chronic TPPU treatment significantly improved cognitive performance, increasing the recognition index to 0.53 ± 0.05 and the discrimination index to 0.05 ± 0.11. These results indicate that TPPU treatment effectively mitigates memory deficits in DM-ADRD rats.

**Figure 3.**
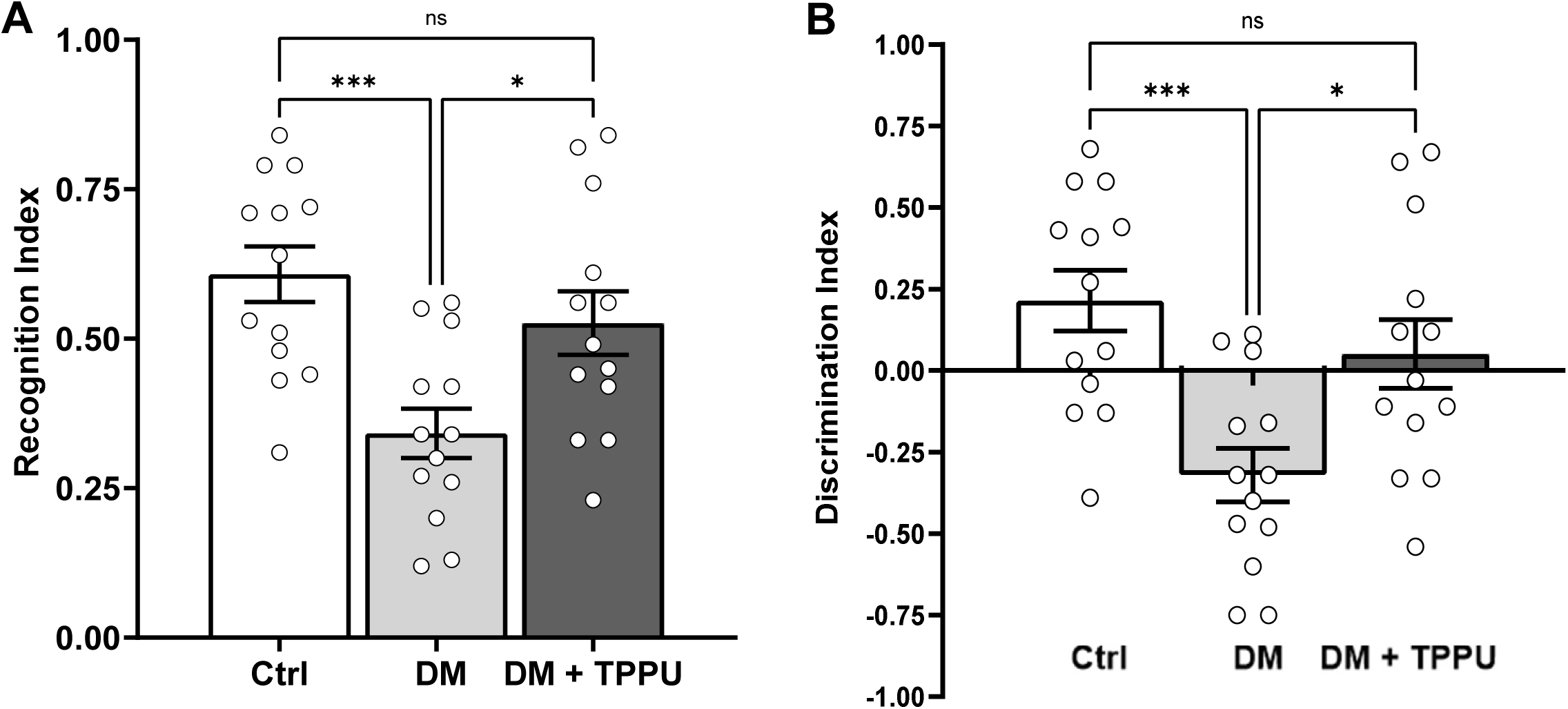
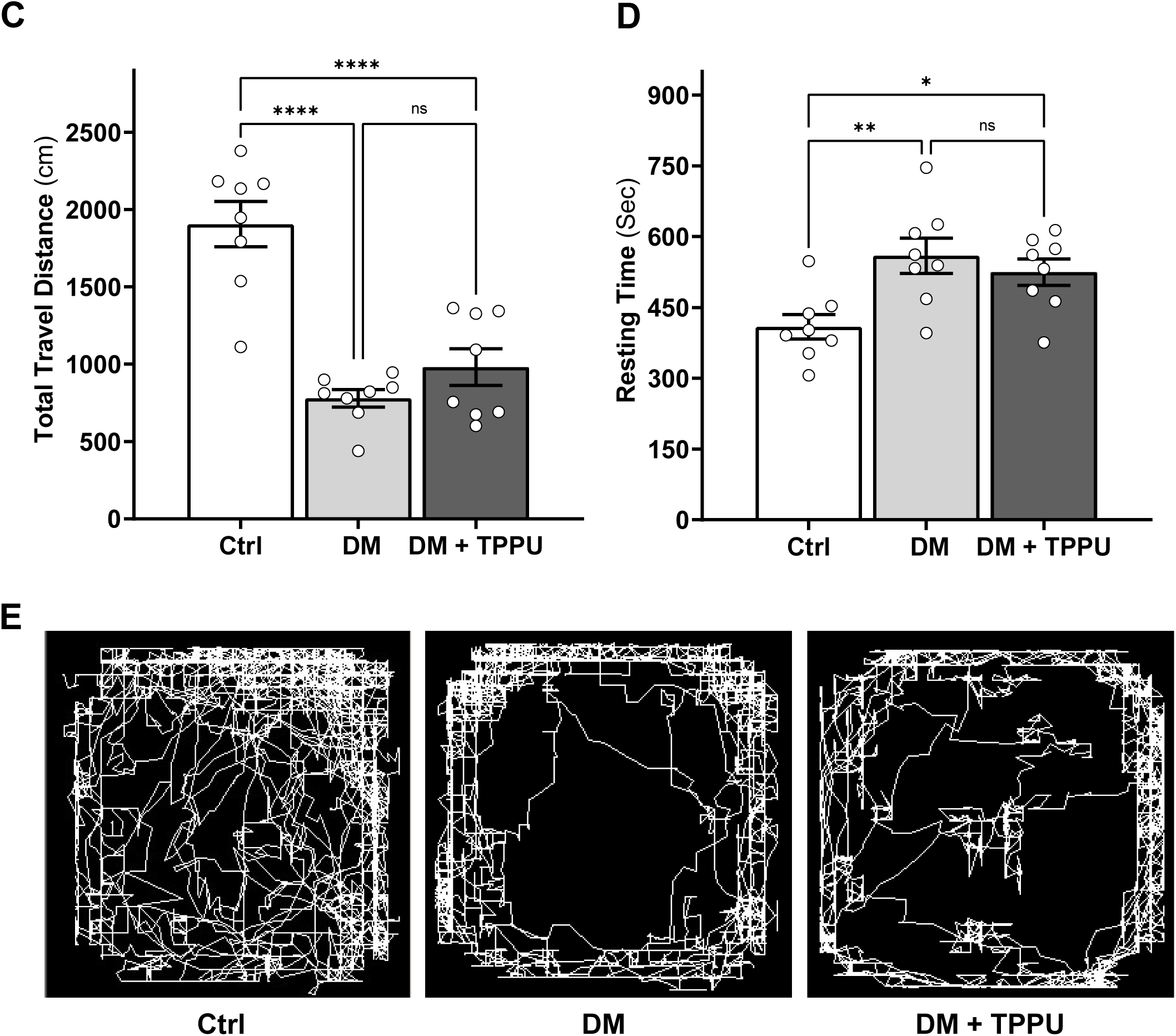
TPPU treatment improves cognitive performance despite persistent locomotor deficits in DM-ADRD rats. **(A)** Recognition Index and **(B)** Discrimination Index from the Novel Object Recognition test. **(C)** Total travel distance and **(D)** resting time from the Open Field Test. **(E)** Representative track maps illustrating exploratory patterns in the open field arena. Data are expressed as mean ± SEM (n = 8-13 rats per group). Statistical significance was determined by one-way ANOVA followed by Tukey’s post-hoc test. \**p* < 0.05, \*\**p* < 0.01, \*\*\**p* < 0.001, \*\*\*\**p* < 0.0001; ns, non-significant.

### Chronic TPPU Treatment Slightly Reduces Anxiety-Like Behavior Without Altering Locomotor Activity in DM-ADRD Rats

General motor activity was assessed using an Open Field Test (**Figures 3C-E**). Compared to controls, DM-ADRD rats showed a significant reduction in total travel distance (779.25 ± 55.91 vs. 1,906.25 ± 146.57 cm) and a corresponding increase in resting time (559.63 ± 37.25 vs. 408.75 ± 25.70 sec). While TPPU treatment resulted in a slight increase in travel distance (981.00 ± 118.28 cm) and a minor decrease in resting time (524.88 ± 27.98 sec), these changes were not statistically significant when compared to the untreated DM group. These findings suggest that while TPPU improves cognitive function, it does not fully restore baseline locomotor activity. However, representative track maps illustrated that while untreated DM rats showed pronounced thigmotaxis, DM + TPPU rats exhibited more frequent forays toward the center of the arena, suggesting that TPPU treatment may slightly reduce anxiety-like behavior in DM-ADRD rats, despite the persistence of overall reduced motor activity.

### Chronic TPPU Treatment Normalizes Transmural Pressure-Dependent Myogenic Responses in Cerebral Vasculature of DM-ADRD Rats

Building on our prior finding that TPPU treatment restores myogenic function in the MCAs and PAs of DM-ADRD rats,^24^ expanded *ex vivo* pressure myography studies were conducted. We assessed both absolute IDs and myogenic tone across a broad range of perfusion pressures, encompassing values both below and above the standard physiological autoregulatory threshold (**Figure 4**). In MCAs, DM-ADRD rats displayed impaired constriction at pressures of 140-180 mmHg, which was successfully reversed by TPPU (**Figure 4A**). Conversely, PAs from DM-ADRD rats exhibited a generalized trend of impaired myogenic reactivity across the pressure spectrum, which was rescued by TPPU administration (**Figure 4B**). We observed that the myogenic tone of the MCA (**Figure 4C**) was elevated in DM-ADRD rats compared with controls at lower perfusion pressures (40 to 100 mmHg), whereas chronic treatment with TPPU significantly increased tone at higher pressures (120 to 180 mmHg), effectively preventing the autoregulatory breakthrough at 120 mmHg observed in untreated DM-ADRD rats.^6,24,44^ In the PA, DM-ADRD rats displayed a higher basal tone at low pressure (10 mmHg) but failed to maintain this constriction at higher pressures (50-60 mmHg). Notably, TPPU reduced the elevated basal tone at lower pressures (10-20 mmHg) while maintaining tone at high pressure (**Figure 4D**). Consistent with our previous reports, these results indicate that although DM-ADRD vessels exhibit higher basal tone at low pressures, they cannot sustain constriction at high pressures.^6,24,44^ Chronic sEH inhibition with TPPU resolves this dysfunction by stabilizing the myogenic response across the autoregulatory spectrum, preventing forced dilation at high pressures, and normalizing basal tone in the PAs.

**Figure 4.**
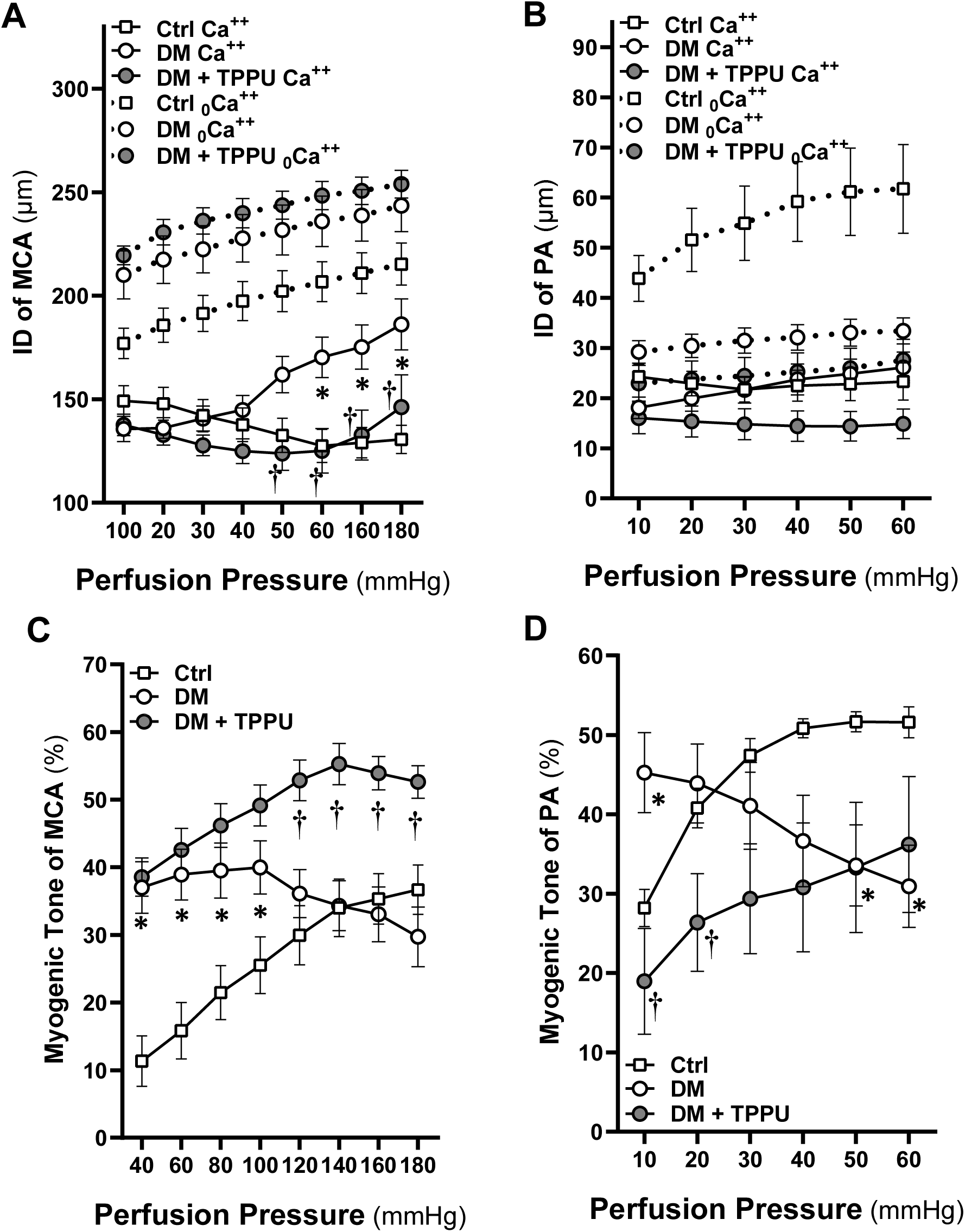
Chronic TPPU treatment restores pressure-induced myogenic reactivity in MCAs and PAs of DM-ADRD rats. **(A)** Internal diameter (ID) of middle cerebral arteries (MCAs) and **(B)** penetrating arterioles (PAs) plotted against incremental perfusion pressures under active and passive conditions. **(C)** The percentage myogenic tone of MCAs across pressures ranging from 40 to 180 mmHg. **(D)** The percentage myogenic tone of PAs across pressures ranging from 10 to 60 mmHg. Data are expressed as mean ± SEM (n = 8 - 12 rats per group). \**p* < 0.05 vs. Control; **†***p* < 0.05 vs. DM-ADRD untreated.

### Chronic TPPU Treatment Rescues Neurovascular Coupling and Restores Capillary-to-Arteriolar Retrograde Signaling in DM-ADRD Rats

As presented in **Figure 5A**, baseline regional cerebral perfusion was significantly reduced in untreated DM-ADRD rats (93.02 ± 2.22%; n = 13) compared to healthy controls (100.30 ± 0.34%; n = 15). Chronic administration of the sEH inhibitor TPPU restored baseline brain perfusion to control levels in DM-ADRD rats (98.98 ± 0.62%; n = 15). Mechanical stimulation of the contralateral whiskers triggered a robust, rapid increase in CBF within the somatosensory barrel cortex of control rats, peaking at a 31.06 ± 1.54% (n = 12) increase from baseline. The functional hyperemia response was profoundly blunted in untreated DM-ADRD rats, showing only a 5.95 ± 1.21% change (n = 6). Chronic TPPU treatment significantly rescued this neurovascular coupling defect, restoring the peak CBF response to 30.06 ± 2.59 % (n = 13), which was indistinguishable from controls (**Figure 5B-C).**

**Figure 5.**
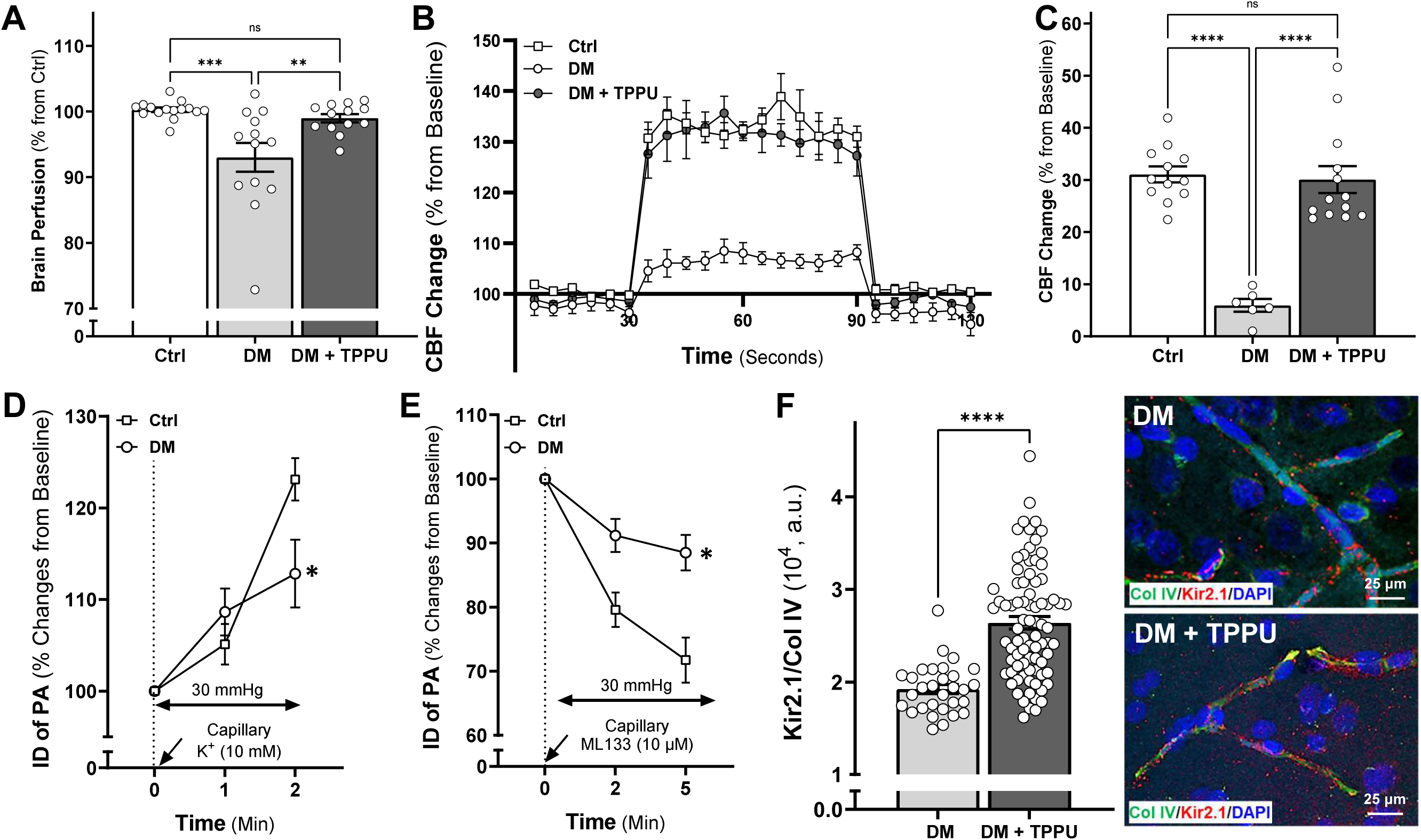
Chronic TPPU treatment restores neurovascular coupling, capillary-to-arteriolar electrical signaling, and microvascular Kir2.1 channel expression in DM-ADRD rats. **(A)** Baseline resting cerebral blood flow (CBF) mapped via laser speckle contrast imaging (LSCI). **(B)** Time-course of changes in cortical CBF during whisker stimulation. **(C)** Average CBF changes during whicker stimulation. **(D)** Upstream arteriolar inner diameter (ID) changes over time following localized capillary application of KCl. **(E)** Upstream arteriolar inner diameter (ID) changes over time following localized capillary application of ML133. **(F)** Representative immunofluorescence images and quantitative analysis of vascular Kir2.1 expression (red) colocalized with Collagen IV (Col IV, green), and nuclei (DAPI, blue). Data are expressed as mean ± SEM (n = 6 - 25 rats per group). Statistical significance was determined by one-way ANOVA followed by Tukey’s post-hoc test for single-point comparisons (A, C, F), and by two-way repeated measures ANOVA followed by Sidak’s post-hoc test for time-course data (B, D, E).

Local micro-application of 10 mM KCl to the distal capillaries in the isolated, pressurized (30 mmHg) CaPA preparations elicited a rapid retrograde hyperpolarization signal, causing a 23.11 ± 2.32 % (n = 25) dilation of the upstream PA in control rats, which was significantly impaired in DM-ADRD rats, yielding only a 12.82 ± 3.69 % (n = 15) dilation (**Figure 5D).** To confirm the functional involvement of inward-rectifier potassium channels, the capillary terminals were exposed to the selective Kir2.1 blocker ML133 (10 μM). In control rats, Kir2.1 blockade rapidly abolished the potassium-induced response and triggered a steep retrograde vasoconstriction, reducing the upstream PA internal diameter to 71.74 ± 3.51 % of baseline (n = 20). Conversely, DM-ADRD vessels exhibited a significantly attenuated constrictor response to ML133 (88.50 ± 2.78 % of baseline; n = 15), demonstrating a loss of functional endothelial Kir2.1 channel activity (**Figure 5E).** Quantitative immunohistochemical analysis of the parenchymal microvasculature (**Figure 5F)** revealed that the colocalization ratio of Kir2.1 channels to the structural basement membrane marker Col IV was significantly higher in TPPU treated DM-ADRD rats [(2.64 ± 0.07) X 10^4^ a.u.] compared to untreated DM-ADRD rats [(1.92 ± 0.05) X 10^4^ a.u.], providing a structural mechanism for the observed functional rescue of retrograde electrical signaling and neurovascular coupling.

### Chronic TPPU Treatment Prevents Blood–Brain Barrier Disruption by Restoring Tight Junction Protein Expression in DM-ADRD Rats

As shown in the representative perfusion maps across a range of transmural pressures (**Figure 6A**), untreated DM rats displayed altered, pressure-dependent microvascular flow distribution with prominent localized "hot spots" of high perfusion at 160 mmHg. This indicates a clear breakthrough in CBF autoregulation, consistent with our previous findings showing that the autoregulatory breakthrough point in DM-ADRD shifts from 140-150 mmHg in control animals to around 120 mmHg. ^6,44^ Remarkably, chronic TPPU treatment effectively stabilized the cortical perfusion profiles and prevented localized perfusion surges at 160 mmHg. Protein expression levels of zonula occludens-1 (ZO-1; **Figure 6B**) and occludin (OCLN; **Figure 6C**) were significantly decreased in the brains of untreated DM rats (0.47 ± 0.06- and 0.50 ± 0.03-fold, respectively) compared to healthy controls (1.00 ± 0.02- and 0.98 ± 0.03-fold, respectively). Remarkably, chronic administration of TPPU rescued this deficit, significantly restoring both ZO-1 and OCLN protein levels (1.13 ± 0.05- and 0.89 ± 0.03-fold increases, respectively) compared with the untreated DM group (**Figure 6D**).

**Figure 6.**
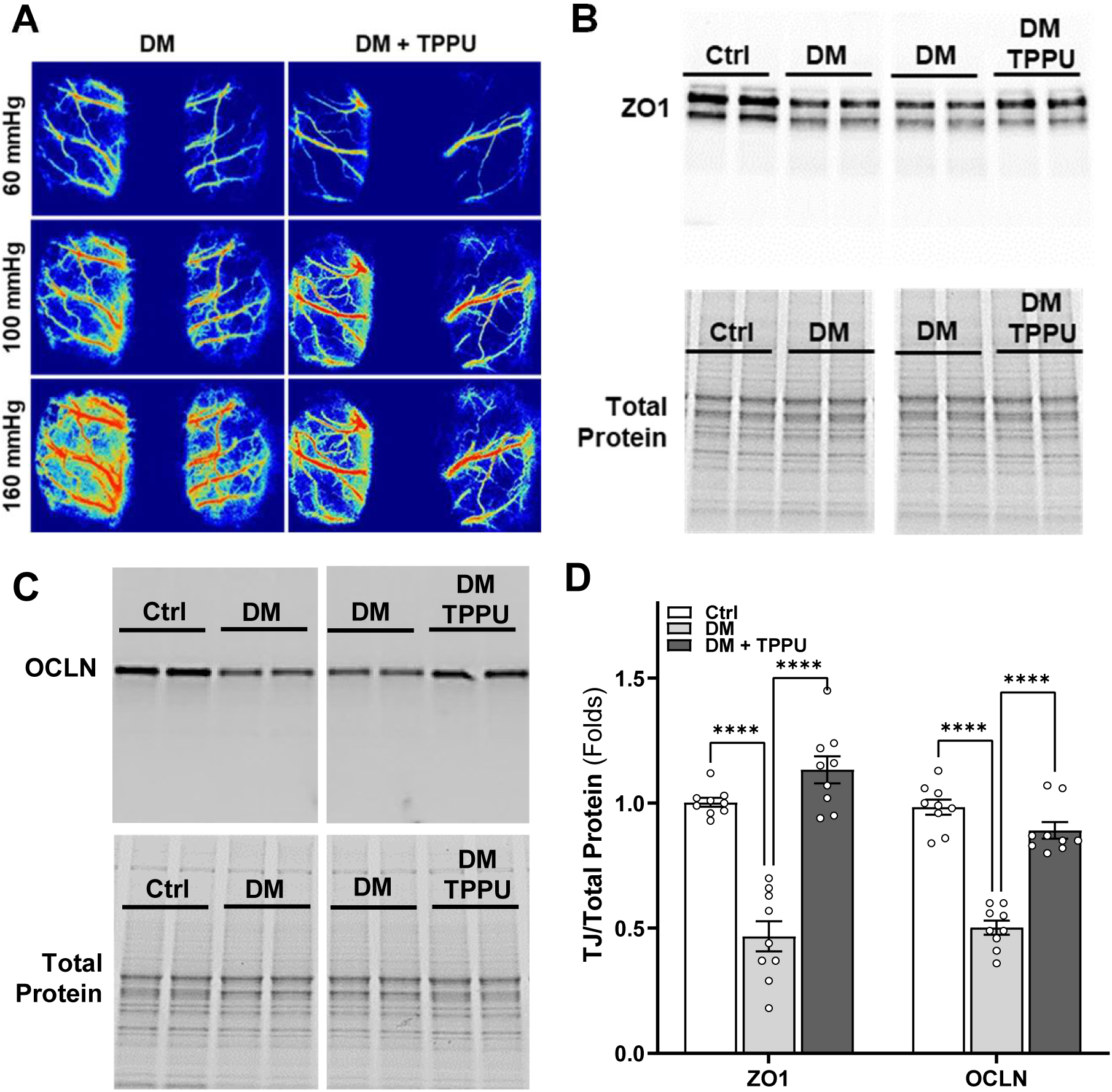
Chronic TPPU treatment prevents blood–brain barrier disruption in DM-ADRD rats. **(A)** Representative laser speckle contrast imaging (LSCI) maps of cortical surface perfusion across a range of mean arterial pressures. (**B-C**) Representative Western blot images and corresponding total protein staining showing cortical protein expression levels of zonula occludens-1 (ZO-1, 200 kD) and occludin (OCLN, 54 kD). **(D)** Quantitative analysis of ZO-1 and OCLN protein expression levels normalized to total protein. Data are presented as mean ± SEM (n = 9 rats per group). Statistical significance was determined using two-way ANOVA followed by Sidak’s post hoc test. \**p* < 0.05, \*\**p* < 0.01, \*\*\**p* < 0.001, \*\*\*\**p* < 0.0001.

### Chronic TPPU Treatment Prevents Microvascular Rarefaction by Restoring Capillary Density in DM-ADRD Rats

As presented in **Figure 7A-B**, capillary density (% area fraction) was significantly decreased in the CA1, CA3, dentate gyrus (DG) of the hippocampus, and cortical (CTX) regions of untreated DM rats (2.93 ± 0.29, 3.48 ± 0.33, 3.06 ± 0.28, and 2.66 ± 0.28%, respectively) compared to control rats (5.59 ± 0.37, 6.31 ± 0.39, 5.15 ± 0.74, and 7.34 ± 0.66%, respectively). Remarkably, chronic administration of TPPU entirely rescued this microvascular deficit, significantly restoring capillary density levels in the hippocampal CA1 (6.94 ± 0.77%), CA3 (6.77 ± 0.60%), DG (5.54 ± 0.33%), and CTX (6.32 ± 0.45%), respectively, suggesting that chronic TPPU treatment effectively preserves microvascular integrity and counteracts capillary rarefaction across multiple vulnerable brain regions in the DM-ADRD model.

**Figure 7.**
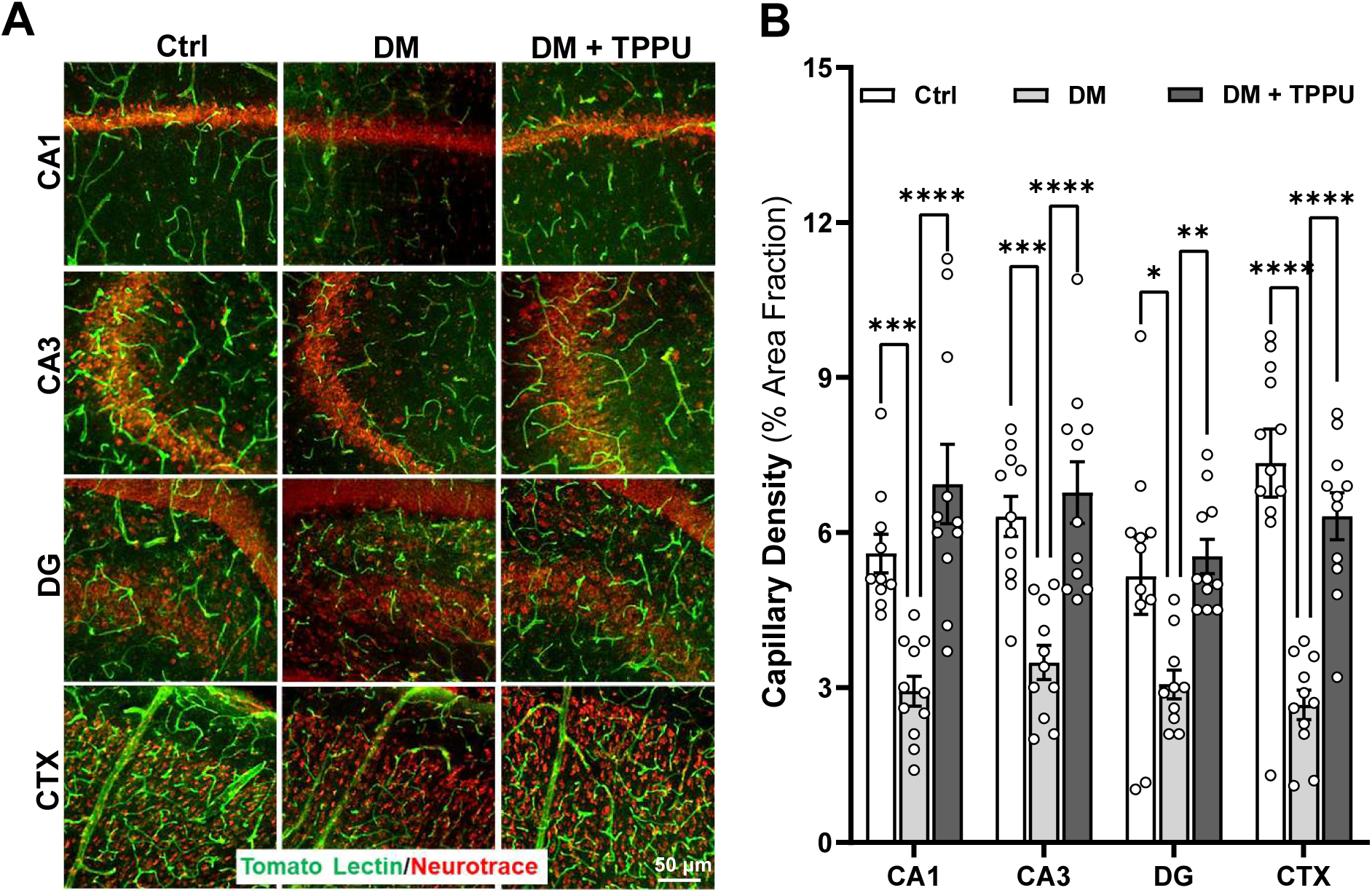
Chronic TPPU treatment restores capillary density across multiple brain regions in DM-ADRD rats. **(A)** Representative immunofluorescence micrographs showing Tomato Lectin-positive capillaries in the CA1, CA3, DG, and CTX regions. **(B)** Quantification of capillary density expressed as a percentage of the total area fraction across the specified brain regions. Data are expressed as mean ± SEM (n = 9 rats per group). Statistical significance was determined using two-way ANOVA followed by Sidak’s post hoc test. \**p* < 0.05, \*\**p* < 0.01, \*\*\**p* < 0.001, \*\*\*\**p* < 0.0001.

### Chronic TPPU Treatment Attenuates Glial Activation in DM-ADRD Rats

As presented in **Figure 8**, hippocampal protein expression of the astrocytic marker GFAP was significantly increased in untreated DM rats (2.11 ± 0.13-fold) compared to control rats (1.02 ± 0.02-fold). Remarkably, chronic administration of TPPU significantly rescued this effect, significantly reducing GFAP expression levels (0.55 ± 0.05-fold; **Figure 8A**). Similarly, protein expression of the microglial marker Iba1 was elevated in untreated DM rats (1.38 ± 0.06-fold) compared to controls (1.02 ± 0.03-fold) and was restored following chronic TPPU treatment (0.95 ± 0.03-fold; **Figure 8B-C**). The number of activated astrocytes was significantly elevated in both the CA1 (30.74 ± 1.46) and CA3 (28.90 ± 1.66) regions of untreated DM rats compared to control rats (4.00 ± 0.59 and 4.39 ± 0.75, respectively; **Figure 8D**). Chronic TPPU treatment significantly attenuated astrogliosis, reducing the counts of activated astrocytes in both the CA1 (17.20 ± 1.25) and CA3 (16.43 ± 1.32) regions of the hippocampus. Similarly, the number of activated microglia was markedly increased in the CA1 (12.26 ± 0.43) and CA3 (14.31 ± 0.86) regions of untreated DM rats compared to control rats (1.46 ± 0.33 and 0.77 ± 0.30, respectively; **Figure 8E**). Chronic TPPU treatment significantly attenuated microgliosis, reducing the counts of activated microglial cells in both the CA1 (6.87 ± 0.83) and CA3 (6.50 ± 0.86) regions. These results suggest that chronic TPPU treatment effectively attenuates astrocytic and microglial activation in the DM-ADRD model.

**Figure 8.**
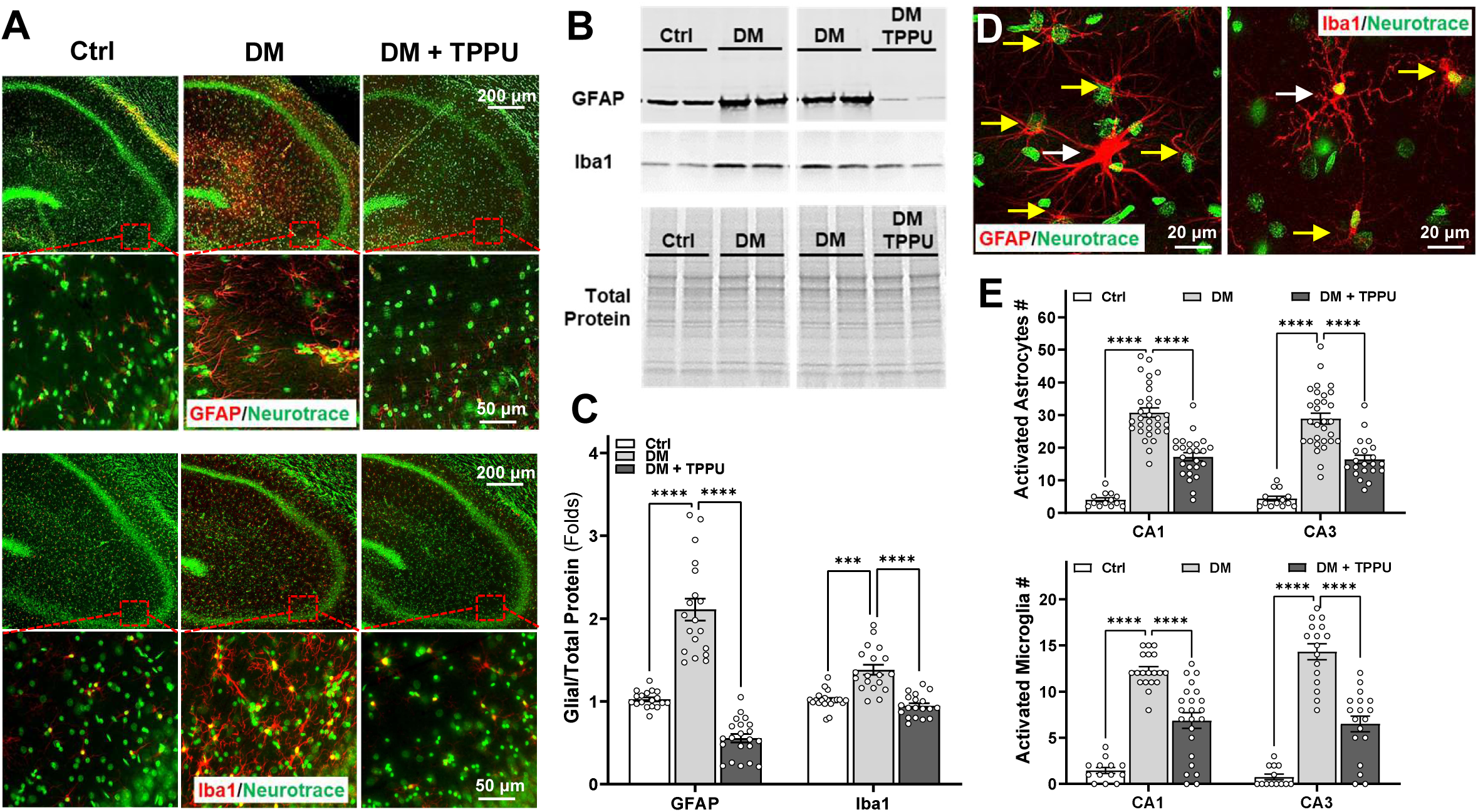
Chronic TPPU treatment attenuates glial activation in DM-ADRD rats. **(A)** Representative immunofluorescence images showing GFAP (astrocytes, red) and Iba1 (microglia, red) co-stained with Neurotrace (green) in the hippocampus. **(B)** Representative Western blot images and corresponding total protein staining for GFAP (50 kD) and Iba1 (17 kD) expression. **(C)** Quantitative analysis of GFAP and Iba1 protein expression levels normalized to total protein. **(D)** High-magnification representative images showing morphological features of activated astrocytes and microglia. **(E)** Quantitative analysis of activated astrocyte and microglia counts in the CA1 and CA3 hippocampal regions. Data are expressed as mean ± SEM (n = 7 - 9 rats per group). Statistical significance was determined using two-way ANOVA followed by Sidak’s post hoc test. \**p* < 0.05, \*\**p* < 0.01, \*\*\**p* < 0.001, \*\*\*\**p* < 0.0001.

### Chronic TPPU Treatment Preserves Neuronal Survival and Synaptic Protein Expression in DM-ADRD Rats

As shown by representative Neurotrace immunofluorescence staining (**Figure 9A**), untreated DM rats exhibited a notable loss of neuronal density and a disruption of cellular architecture within the hippocampal subfields. Quantitative analysis of neuron counts (**Figure 9B**) confirmed a significant reduction in neuronal numbers within both the CA1 (96.13 ± 2.28) and CA3 (85.44 ± 2.68) regions of untreated DM rats compared to healthy controls (111.94 ± 1.26 and 97.44 ± 2.01, respectively). Remarkably, chronic administration of TPPU significantly prevented this neurodegenerative progression, restoring neuronal survival in both the CA1 (104.13 ± 2.23) and CA3 (94.79 ± 3.34) regions. To evaluate synaptic integrity, we assessed the expression of the postsynaptic PSD95 and the presynaptic Synaptophysin (SY38) proteins via Western blotting (**Figure 9C**). Quantitative analysis (**Figure 9D**) demonstrated a significant downregulation of both PSD95 (0.42 ± 0.07) and SY38 (0.74 ± 0.02) protein levels (expressed as fold change) in the hippocampus of untreated DM rats compared to controls. This loss of synaptic markers was rescued by chronic TPPU treatment, which significantly increased expression levels of both PSD95 (0.71 ± 0.08) and SY38 (0.97 ± 0.05), suggesting that chronic TPPU treatment effectively mitigates neurodegeneration and maintains synaptic architecture in the DM-ADRD model.

**Figure 9.**
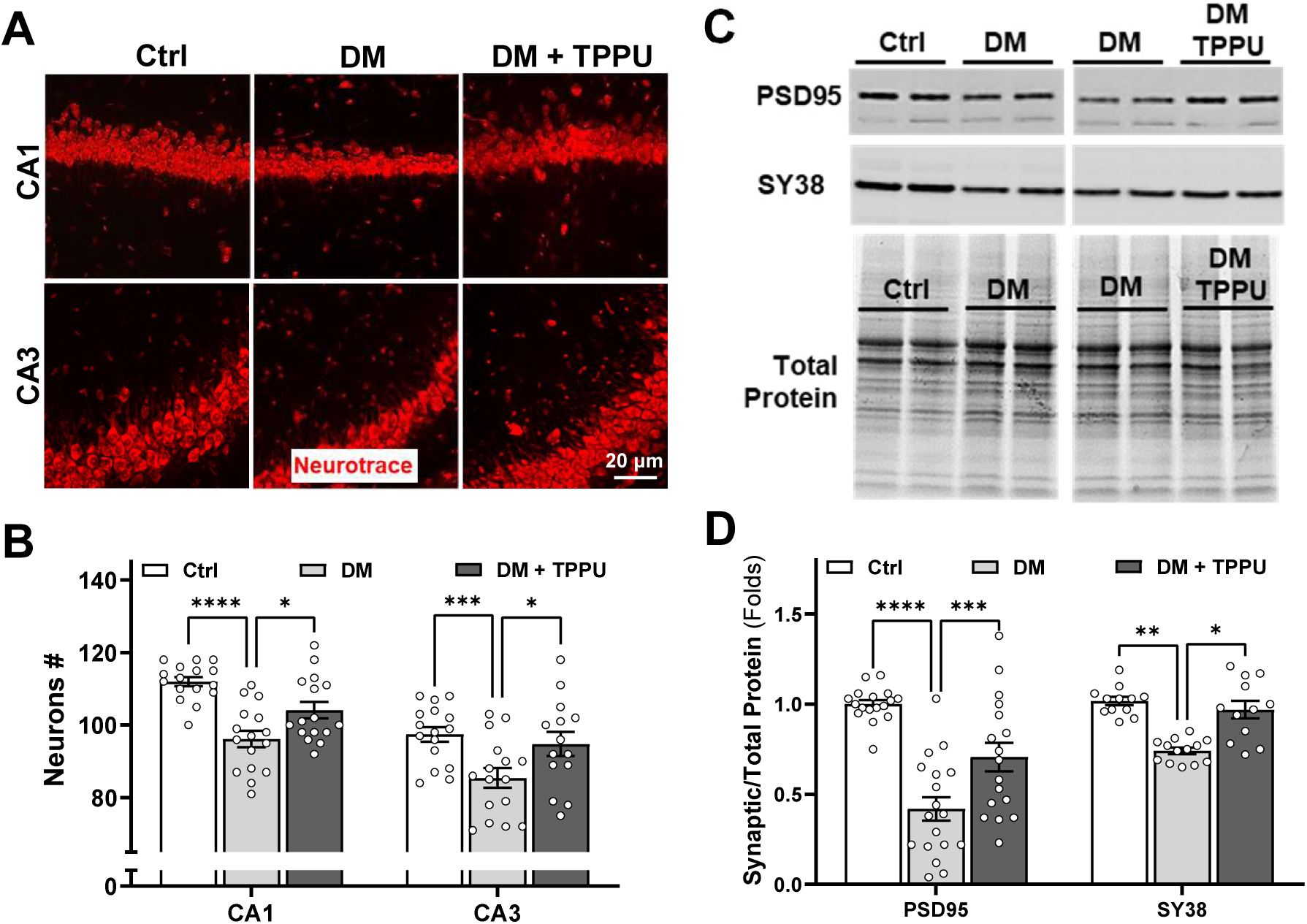
Chronic TPPU treatment preserves neuronal survival and synaptic protein expression in DM-ADRD rats. **(A)** Representative immunofluorescence images of Neurotrace-stained neurons in the CA1 and CA3 hippocampal regions. **(B)** Quantitative analysis of neuron counts within the CA1 and CA3 regions. **(C)** Representative Western blot images and corresponding total protein expression for PSD95 (95 kD) and Synaptophysin (SY38, 38 kD) expression in the brain. **(D)** Quantitative analysis of hippocampal PSD95 and SY38 expression levels normalized to total protein. Data are expressed as mean ± SEM (n = 7 - 9 rats per group). Statistical significance was determined using two-way ANOVA followed by Sidak’s post hoc test\**p* < 0.05, \*\**p* < 0.01, \*\*\**p* < 0.001, \*\*\*\**p* < 0.0001.

### Chronic TPPU Treatment Selectively Regulates Brain Chemokine Levels Without Altering Baseline Pro-Inflammatory Cytokines in DM-ADRD Rats

To determine the impact of chronic sEH inhibition on central inflammatory signaling, we profiled the expression levels of key inflammatory cytokines and chemokines via multiplex ELISA. As shown in **Figure 10A**, brain levels of classic pro-inflammatory cytokines, including tumor necrosis factor-alpha (TNFα), interleukin-6 (IL-6), interferon-gamma (IFNγ), interleukin-1 alpha (IL-1α), and interleukin-1 beta (IL-1β), remained statistically unaltered among all groups. In contrast, quantitative analysis of chemokine levels (**Figure 10B**) revealed a profound upregulation in the untreated DM-ADRD group, with elevated expression levels of monocyte chemoattractant protein-1 (MCP-1; 1.29 ± 0.04-fold), RANTES (CCL5, 1.40 ± 0.08-fold), and macrophage inflammatory protein-1 alpha (MIP-1α; 1.50 ± 0.09-fold), respectively, compared to control animals. Chronic TPPU treatment prevented this increase, significantly suppressing MCP-1 (1.02 ± 0.03-fold), RANTES (0.97 ± 0.04-fold), and MIP-1α (0.81 ± 0.03-fold), respectively. These results suggest that chronic TPPU treatment selectively dampens specific chemokine signaling networks in the DM-ADRD model.

**Figure 10.**
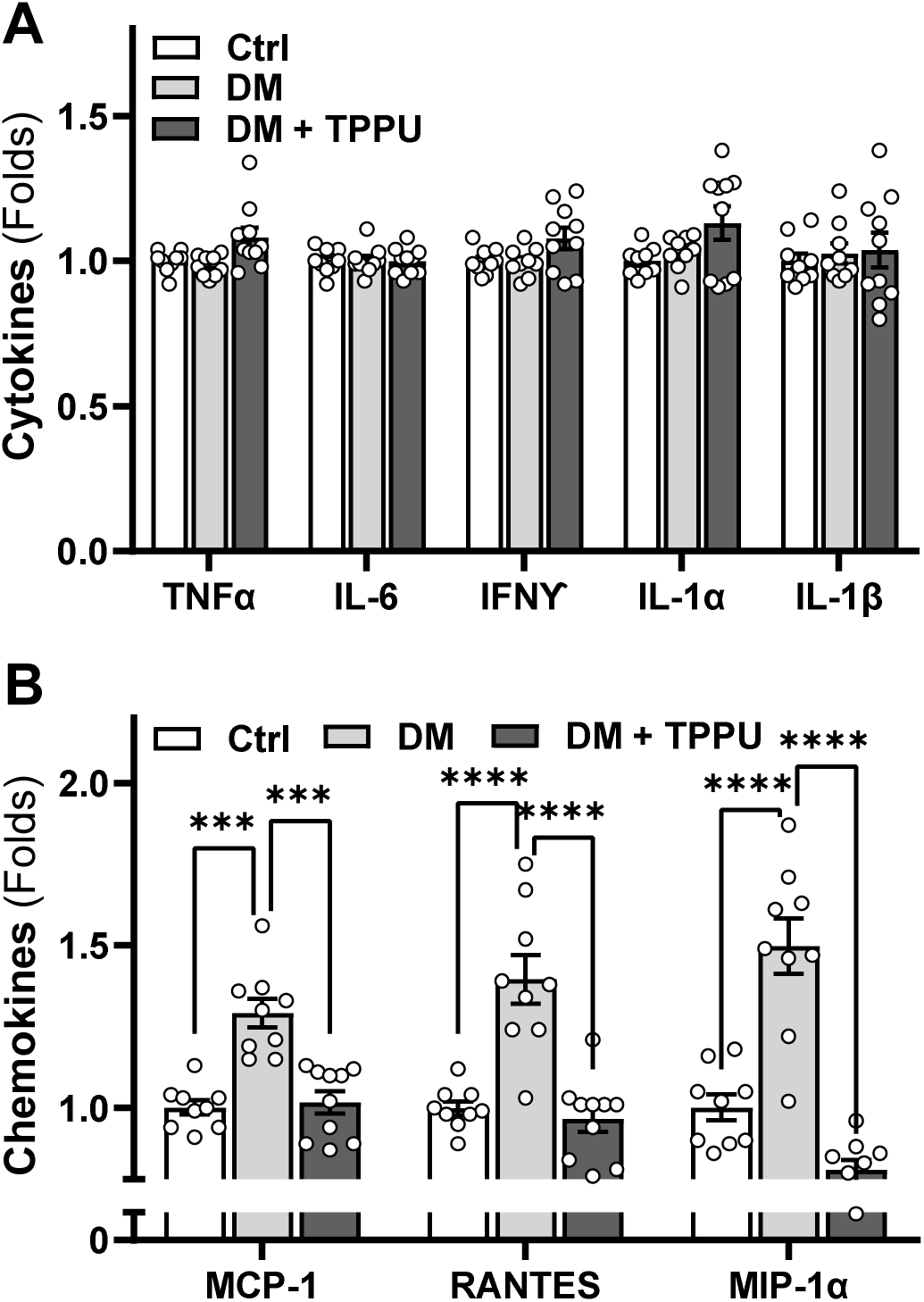
Chronic TPPU treatment selectively regulates brain chemokine levels without altering baseline pro-inflammatory cytokines in DM-ADRD rats. **(A)** Quantitative analysis of cortical pro-inflammatory cytokine expression levels. **(B)** Quantitative analysis of cortical chemokine expression levels. Data are expressed as mean ± SEM (n = 8 rats per group). Statistical significance was determined using two-way ANOVA followed by Sidak’s post hoc test\**p* < 0.05, \*\**p* < 0.01, \*\*\**p* < 0.001, \*\*\*\**p* < 0.0001.

## DISCUSSION

Compounding the global threat of cognitive decline, DM, especially type 2 DM, is a progressive metabolic disorder that significantly increases the risk of ADRD.^2^ As a global health crisis affecting 14% of adults, totaling more than 830 million people worldwide,^58^ untreated or poorly controlled DM drives a multifactorial deterioration of cerebrovascular and cognitive function for which there are currently no curative or effective preventive therapies to stop or slow progression. In the disease state, chronic hyperglycemia, insulin resistance, and other pathological cascades, such as accelerated advanced glycation end-product accumulation, chronic low-grade inflammation, and elevated oxidative stress, are thought to trigger severe cerebrovascular pathology.^6^ Accumulating preclinical and clinical evidence indicates that this damage is driven by a multifaceted deficit of vascular and glymphatic homeostatic mechanisms, which often precedes classic neurodegenerative pathologies and cognitive symptoms.^5,24,59,60^ This breakdown is characterized by a reduction in baseline CBF, impairment of myogenic autoregulation via VSMC or pericyte dysfunction, loss of neurovascular coupling and functional hyperemic responses, structural decay of TJs at the BBB, and compromised glymphatic clearance linked to astrocytic endfeet depolarization.^6,12,14,17^ Concurrently, genome-wide association studies and molecular profiling have identified the genetic upregulation of sEH as a critical biochemical contributor to this microvascular vulnerability.^23-25^ Because sEH rapidly converts highly protective, anti-inflammatory, and vasodilatory EETs and other epoxyfatty acids into less bioactive diols, elevated sEH levels in the diabetic brain accelerate microvascular degradation. ^24,45,46^ This enzymatic shift drives a self-reinforcing cycle of chronic neuroinflammation, capillary stalling, and synaptic injury.^47,48^ Therefore, establishing a clear understanding of these untreated microvascular pathologies provides a necessary benchmark for evaluating therapeutic strategies, shifting paradigms beyond neuron-centric models.

We have previously reported that targeting systemic pathways with the SGLT2 inhibitor luseogliflozin attenuated mitochondrial dysfunction and oxidative stress in cerebral VSMCs, restored CBF autoregulation and functional hyperemia, reduced BBB leakage, and improved cognitive performance across both AD and DM-ADRD rat models.^43,44^ Notably, while this SGLT2 inhibition drove robust systemic metabolic effects (reducing body weight, plasma glucose, and HbA1c levels toward the normoglycemic range) exclusively in the DM-ADRD model, it rescued neurovascular and cognitive function in the non-diabetic AD model without inducing any systemic metabolic alterations, demonstrating that luseogliflozin can improve cerebral vascular function both by lowering glucose or through glucose-independent mechanisms. Parallel to the SGLT2 inhibition studies, we found that sEH protein expression is upregulated in the brain of the AD rat model and that systemic sEH inhibition via TPPU similarly improves cerebral hemodynamics and hippocampus-based spatial learning and memory in both AD and DM-ADRD models.^24^ Intriguingly, while chronic TPPU treatment did not alter body weight in either model, it significantly reduced plasma glucose and HbA1c levels only in the DM-ADRD group; however, unlike SGLT2 inhibition, glycemia remained firmly within the diabetic range and was not normalized.^24^ Transcriptomic and molecular profiling from these prior investigations in AD rats revealed that TPPU protects this vulnerable environment by directly suppressing vascular inflammation, oxidative stress, Aβ plaque burden, BBB leakage, and glial activation.^24,38^ The protective effects of sEH inhibition were further amplified when pairing sEH with COX-2 dual inhibition by simultaneously blocking the conversion of EETs and preventing arachidonic acid metabolism into pro-inflammatory prostaglandins. Dual inhibition acts cooperatively to blunt chronic neurovascular inflammation and robustly attenuate aberrant vascular remodeling.^39^

The present study builds on this established foundation by demonstrating that chronic administration of TPPU (1 mg/kg/day) downregulates brain sEH protein expression, rescues cortical- and hippocampal-mediated long-term non-spatial recognition memory as measured by the NOR test, and reduces anxiety in DM-ADRD rats. We reveal that the stabilization of the myogenic response and tone in MCAs and PAs, alongside the normalization of impaired whisker-evoked functional hyperemia, directly drives these improvements in brain perfusion via a novel upregulation of capillary Kir2.1 expression. At the structural level, this targeted intervention restores the essential TJ proteins ZO-1 and OCLN, mitigates capillary rarefaction, and suppresses astrocyte and microglial activation. At the cellular level, TPPU attenuates hippocampal neurodegeneration, preserves synaptic integrity by restoring the expression of PSD95 and SY38, and dampens the local microvascular inflammatory cascade by profoundly reducing key pro-inflammatory chemokines, including MCP-1, RANTES, and MIP-1α.

Our previous work demonstrated that TPPU reverses spatial learning and memory deficits in DM-ADRD rats using an eight-arm water maze paradigm.^24^ In the current study, we first validated this finding and then broadened our behavioral characterization by utilizing the NOR and Open Field Tests to assess anxiety-like behaviors alongside long-term non-spatial recognition memory processed via interconnected cortical and hippocampal loops. Introducing the Open Field Test enabled us to explicitly evaluate whether advanced age, when paired with type 2 diabetes mellitus, induces intrinsic locomotor or exploratory impairments. We found that despite sEH inhibition significantly rescuing non-spatial recognition memory, as shown by a restored discrimination index during the NOR testing phase following a four-hour retention interval, it did not alter DM-related intrinsic locomotor or exploratory impairments. Furthermore, TPPU-treated DM-ADRD rats attenuated anxiety-like phenotypes, spending significantly more time in the center zone and traveling greater total distances in the open arena. These outcomes illuminate the age-dependent progression of DM-ADRD. While young diabetic rats exhibit isolated spatial memory deficits, as we previously reported,^6^ advanced age leads to an extensive decline in both spatial and non-spatial memory domains. This progressive divergence aligns with classic lesion studies showing that minor hippocampal damage suffices to impair spatial navigation, whereas non-spatial recognition depends on expansive cortical networks and requires greater neuronal dysfunction and structural loss before collapsing. By rescuing non-spatial recognition memory in aged DM-ADRD rats, TPPU effectively preserves the functional connectivity of the perirhinal, entorhinal, and prefrontal cortices, as well as their hippocampal relays. Furthermore, given that diabetic states frequently provoke anxiety linked to elevated pro-inflammatory cytokines, the successful mitigation of anxiety-like traits indicates that stabilizing the neurovascular unit via sEH inhibition counteracts the chronic microvascular stress and activation of chemokines that typically disrupt emotional processing within frontocortical and amygdalar networks.

Uncontrolled fluctuations in CBF play a direct role in driving BBB disintegration and capillary rarefaction within the diabetic brain. When the protective mechanism of myogenic response and autoregulation fails, elevated systemic blood pressure is transmitted into the delicate microvasculature. This hemodynamic stress generates high-pressure surges and aberrant flow distribution that distend and damage the endothelium, leading to the structural decay of critical TJ assemblies and subsequent BBB leakage. Over time, this chronic mechanical trauma and sustained hypoperfusion trigger capillary regression, causing severe microvascular rarefaction that starves metabolic hotspots of necessary oxygen and nutrients.^4,13,61,62^ Our present findings demonstrate that chronic sEH inhibition with TPPU counteracts loss of capillary and cerebral hypoperfusion. This therapeutic efficacy is underpinned by the optimized pharmacokinetics and favorable physicochemical properties of TPPU, including a low molecular weight of 359.3 g/mol, a low topological polar surface area, and ideal lipophilicity (log *P* ≈ 2.5).^54,63^ These attributes confer this small-molecule inhibitor, excellent BBB permeability, resulting in highly effective accumulation in brain tissue and robust target engagement under chronic administration.^64,65^ By efficiently penetrating the central nervous system to stabilize cortical perfusion profiles and prevent high-pressure breakthrough surges, TPPU shields the microvasculature from mechanical trauma. This preservation enables restoration of ZO-1 and OCLN protein expression and rescues capillary density in highly vulnerable CTX and hippocampal subregions, including CA1, CA3, and DG.

Downstream of these vascular structural rescues, chronic TPPU administration effectively dampens cerebral inflammation by directly suppressing the chemokines driving glial activation and glymphatic failure. Microvascular endothelial injury triggers a localized inflammatory cascade marked by the aberrant release of the chemo-attractants MCP-1, RANTES, and MIP-1α, which specifically recruit and transform resting microglia and astrocytes into highly reactive, pro-inflammatory phenotypes. This pathological glial activation forces astrocytic endfeet depolarization and the loss of polarized AQP4 water channels, disrupting glymphatic clearance networks and trapping neurotoxic metabolites in the parenchyma. Our findings demonstrate that standalone sEH inhibition profoundly reduces microvascular MCP-1, RANTES, and MIP-1α expressions, directly correlating with the suppression of reactive gliosis. Intriguingly, broad systemic cytokines remain unaltered because TPPU does not normalize hyperglycemia, indicating that systemic metabolic inflammation persists; however, by stabilizing EETs, TPPU selectively protects the local endothelial wall, blocking focal chemokine transcription and glial recruitment without altering systemic immune status. This focal microvascular protection stems from the ability of preserved EETs to directly inhibit nuclear factor-kappa B translocation into the nucleus within the endothelium, thereby silencing downstream leukocyte-adhesion signatures and focal chemoattractant signals.^66^ By mitigating this chemokine-driven glial activation, TPPU may preserve AQP4 polarization to restore glymphatic drainage while successfully breaking the self-reinforcing loop of neuroinflammation and capillary stalling.^67-70^ Chronic neuroinflammation and sustained hypoperfusion in the untreated DM-ADRD brain routinely trigger synaptic pruning and neuronal apoptosis, marked by catastrophic loss of essential pre- and postsynaptic scaffolding elements.^38^ By simultaneously arresting microvascular leakage, reactive gliosis, and glymphatic clearance failure, chronic TPPU administration establishes a protected parenchymal environment that preserves pre- and post- synaptic densities and opposes progressive neurodegeneration.

In summary, sEH inhibition delivers neurovascular, glymphatic, and synaptic protection, establishing a therapeutic approach to interrupt the progressive pathological cascades of DM-ADRD without requiring the normalization of systemic metabolism. Several limitations must be considered, including our exclusive focus on a male rodent cohort, which necessitates future validation in female models to account for known sexual dimorphism. Additionally, the precise downstream intracellular signaling pathways connecting microvascular EET stabilization to localized chemokine suppression require further molecular dissection, and safety profiles under prolonged administration conditions remain to be characterized. Taken together, these data underscore that TPPU preserves cognitive function in DM-ADRD by directly counteracting neuroinflammation, gliosis, and cerebrovascular dysfunction while protecting neuronal survival and synaptic machinery, establishing sEH inhibition as a powerful strategy that delivers comprehensive neurovascular rescue even in the presence of sustained, systemic diabetic hyperglycemia. Nevertheless, the present study provided definitive proof of concept that targeting microvascular stability can limit structural neurodegeneration and cognitive decline in DM-ADRD, offering a paradigm-shifting therapeutic strategy that protects the brain even with sustained systemic hyperglycemia. Targeting sEH represents an integrated disease-modifying strategy to restore cerebrovascular integrity, stabilize immune homeostasis, and arrest the progression of cognitive deficits in DM-ADRD.

## DECLARATIONS

### Ethics approval and consent to participate

Not applicable.

### Conflict of Interest

B.D. Hammock is a founder; S.H. Hwang and J. Yang are part-time employees of EicOsis L.L.C., a startup company with an sEH inhibitor in human clinical trials. B.D. Hammock, S.H., Hwang, C., Morisseau, and J. Yang are inventors on patents owned by the University of California for the design and pharmaceutical use of sEH inhibitors.

### Author Contributions

F.F. and R.J.R. conceived and designed research; X.F., J.J.B., H.Z., G.C.M., A.G., H.Y., and S.H.H. performed experiments; X.F., J.J.B., H.Z., G.C.M., A.G., F.F., and R.J.R. analyzed data; X.F., J.J.B., H.Z., G.C.M., A.G., H.Y., R. D., J.Y., S.H.H., C. M., B.D.H., R.J.R., and F.F. interpreted results; X.F. and F.F. prepared figures; X.F. and F.F. drafted the manuscript; all authors edited, revised, and approved the final version of the manuscript.

### Consent for publication

All participants have consented to the publication.

### Funding

This study was supported by AG079336, AG057842, AG094068, R35ES030443, and U54NS127758 from the National Institutes of Health; A25-1690 from the Harrington Discovery Institute; 25PRE1365157 and 25POST1363814 from the American Heart Association; and TRIBA/ Physiology Faculty Startup Fund from Augusta University.

### Data availability

The data supporting the findings of this study are available from the corresponding author upon reasonable request.

## REFERENCES

1. Agrawal M, Agrawal AK. Pathophysiological Association Between Diabetes Mellitus and Alzheimer’s Disease. Cureus. 2022;14:e29120. doi: 10.7759/cureus.29120

2. 2025 Alzheimer’s disease facts and figures. Alzheimers Dement. 2025;29:e70235. doi: 10.1002/alz.70235

3. Wu J, Li J, Qin X, Chen W. Association between diabetes mellitus and risk of Alzheimer’s disease: a meta-analysis and systematic review. Front Endocrinol (Lausanne). 2026;17:1736410. doi: 10.3389/fendo.2026.1736410

4. Fan F, Roman RJ. Reversal of cerebral hypoperfusion: a novel therapeutic target for the treatment of AD/ADRD? Geroscience. 2021;43:1065–1067. doi: 10.1007/s11357-021-00357-7

5. Fang X, Tang C, Zhang H, Border JJ, Liu Y, Shin SM, Yu H, Roman RJ, Fan F. Longitudinal characterization of cerebral hemodynamics in the TgF344-AD rat model of Alzheimer’s disease. Geroscience. 2023;45:1471–1490. doi: 10.1007/s11357-023-00773-x

6. Wang S, Lv W, Zhang H, Liu Y, Li L, Jefferson JR, Guo Y, Li M, Gao W, Fang X, et al. Aging exacerbates impairments of cerebral blood flow autoregulation and cognition in diabetic rats. Geroscience. 2020;42:1387–1410. doi: 10.1007/s11357-020-00233-w

7. Anwer M, Poliakova T, Albanus RD, Bhuiyan MIH, Brickman AM, Chaudhuri S, Cuddy L, Eisenbaum M, Foley K, Gould DB, et al. Vascular contribution to cognitive impairment and dementia (VCID): proceedings of 2025 workshop of the Jackson Laboratory. Mamm Genome. 2026;37. doi: 10.1007/s00335-026-10210-x

8. Kang SH, Kim S, Kim YJ, Yoo H, Lee EH, Jang H, Shin D, Yun J, Kim JP, Kim HJ, et al. Glymphatic dysfunction links vascular pathology to Alzheimer’s biomarkers and cognitive decline. Alzheimers Res Ther. 2026. doi: 10.1186/s13195-026-01964-2

9. Wang S, Tang C, Liu Y, Border JJ, Roman RJ, Fan F. Impact of impaired cerebral blood flow autoregulation on cognitive impairment. Front Aging. 2022;3:1077302. doi: 10.3389/fragi.2022.1077302

10. Fan F, Zhao N, Guo M. Lymphatic-venous anastomosis: Cracking the code of Alzheimer’s disease treatment? Neural Regen Res. 2025:doi: 10.4103/NRR.NRR–D–4125–00540. . doi: 10.4103/NRR.NRR-D-25-00540

11. Fan F, Geurts AM, Murphy SR, Pabbidi MR, Jacob HJ, Roman RJ. Impaired myogenic response and autoregulation of cerebral blood flow is rescued in CYP4A1 transgenic Dahl salt-sensitive rat. Am J Physiol Regul Integr Comp Physiol. 2015;308:R379–390. doi: 10.1152/ajpregu.00256.2014

12. Liu Y, Wang S, Guo Y, Zhang H, Roman RJ, Fan F. Impaired Pericyte Constriction and Cerebral Blood Flow Autoregulationin Diabetes. Stroke. 2020;51:AWP498–AWP498.

13. Shekhar S, Wang S, Mims PN, Gonzalez-Fernandez E, Zhang C, He X, Liu CY, Lv W, Wang Y, Huang J, et al. Impaired Cerebral Autoregulation-A Common Neurovascular Pathway in Diabetes may Play a Critical Role in Diabetes-Related Alzheimer’s Disease. Curr Res Diabetes Obes J. 2017;2:555587.

14. Guo Y, Wang S, Liu Y, Fan L, Booz GW, Roman RJ, Chen Z, Fan F. Accelerated cerebral vascular injury in diabetes is associated with vascular smooth muscle cell dysfunction. Geroscience. 2020;42:547–561. doi: 10.1007/s11357-020-00179-z

15. Fernandez-Klett F, Offenhauser N, Dirnagl U, Priller J, Lindauer U. Pericytes in capillaries are contractile in vivo, but arterioles mediate functional hyperemia in the mouse brain. Proc Natl Acad Sci U S A. 2010;107:22290–22295. doi: 10.1073/pnas.1011321108

16. Harder DR, Alkayed NJ, Lange AR, Gebremedhin D, Roman RJ. Functional hyperemia in the brain: hypothesis for astrocyte-derived vasodilator metabolites. Stroke. 1998;29:229–234. doi: 10.1161/01.str.29.1.229

17. Liu Y, Zhang H, Wang S, Guo Y, Fang X, Zheng B, Gao W, Yu H, Chen Z, Roman RJ, et al. Reduced pericyte and tight junction coverage in old diabetic rats are associated with hyperglycemia-induced cerebrovascular pericyte dysfunction. Am J Physiol Heart Circ Physiol. 2021;320:H549–H562. doi: 10.1152/ajpheart.00726.2020

18. Fang X, Border JJ, Rivers PL, Zhang H, Williams JM, Fan F, Roman RJ. Amyloid beta accumulation in TgF344-AD rats is associated with reduced cerebral capillary endothelial Kir2.1 expression and neurovascular uncoupling. Geroscience. 2023;45:2909–2926. doi: 10.1007/s11357-023-00841-2

19. Crumpler R, Roman RJ, Fan F. Capillary Stalling: A Mechanism of Decreased Cerebral Blood Flow in AD/ADRD. J Exp Neurol. 2021;2:149–153. doi: 10.33696/neurol.2.048

20. Yoon J-H, Shin P, Joo J, Kim GS, Oh W-Y, Jeong Y. Increased capillary stalling is associated with endothelial glycocalyx loss in subcortical vascular dementia. Journal of Cerebral Blood Flow & Metabolism. 2022;42:1383–1397. doi: 10.1177/0271678x221076568

21. Simon M, Wang MX, Ismail O, Braun M, Schindler AG, Reemmer J, Wang Z, Haveliwala MA, O’Boyle RP, Han WY, et al. Loss of perivascular aquaporin-4 localization impairs glymphatic exchange and promotes amyloid β plaque formation in mice. Alzheimers Res Ther. 2022;14:59. doi: 10.1186/s13195-022-00999-5

22. Xue H, Bi S, Chen Z, He Y, Xu W, Xu X, Cui B, Hu Y, Qi Z, Yan S, et al. Glymphatic system dysfunction mediates amyloid deposition and cognitive impairment in Alzheimer’s disease: a PET/MRI multimodality imaging study. EJNMMI Res. 2025;16:2. doi: 10.1186/s13550-025-01339-y

23. Cheng F, Hou Y, Zhang P, Fan F, Haines JL, Pieper AA, McReynolds C, Hammock BD, Leverenz JB, Cummings JL. Alzheimer’s disease genetics and real-world patient data fuel target and drug discovery in European and African American Ancestries using AI/ML approaches. Alzheimer’s & Dementia. 2025;20:e087887.

24. Tang C, Border JJ, Zhang H, Gregory A, Bai S, Fang X, Liu Y, Wang S, Hwang SH, Gao W, et al. Inhibition of soluble epoxide hydrolase ameliorates cerebral blood flow autoregulation and cognition in alzheimer’s disease and diabetes-related dementia rat models. Geroscience. 2025;47:4429–4449. doi: 10.1007/s11357-025-01550-8

25. Cheng F, Hou Y, Zhang P, Li Y, Lorincz-Comi N, Fan F, Song W, Fang X, Qiu Y, Paz IR, et al. Combining genetics with real-world patient data enables ancestry-specific target identification and drug discovery in Alzheimer’s disease. Research Square (Preprint). 2024. doi: 10.21203/rs.3.rs-5716817/v1

26. Ma SL, Tang NL, Zhang YP, Ji LD, Tam CW, Lui VW, Chiu HF, Lam LC. Association of prostaglandin-endoperoxide synthase 2 (PTGS2) polymorphisms and Alzheimer’s disease in Chinese. Neurobiol Aging. 2008;29:856–860. doi: 10.1016/j.neurobiolaging.2006.12.011

27. Fan F, Simino J, Auchus AP, Knopman DS, Boerwinkle E, Fornage M, Mosley TH, Roman RJ. Functional variants in CYP4A11 and CYP4F2 are associated with cognitive impairment and related dementia endophenotypes in the elderly. In: The 16th International Winter Eicosanoid Conference. Baltimore; 2016:CV5.

28. Fan F, Ge Y, Lv W, Elliott MR, Muroya Y, Hirata T, Booz GW, Roman RJ. Molecular mechanisms and cell signaling of 20-hydroxyeicosatetraenoic acid in vascular pathophysiology. Front Biosci (Landmark Ed). 2016;21:1427–1463.

29. Roman RJ, Fan F. 20-HETE: Hypertension and Beyond. Hypertension. 2018;72:12–18. doi: 10.1161/HYPERTENSIONAHA.118.10269

30. Gonzalez-Fernandez E, Fan L, Wang S, Liu Y, Gao W, Thomas KN, Fan F, Roman RJ. The adducin saga: Pleiotropic genomic targets for precision medicine in human hypertension; vascular, renal, and cognitive diseases. Physiol Genomics. 2021. doi: 10.1152/physiolgenomics.00119.2021

31. Fan F, Pabbidi MR, Ge Y, Li L, Wang S, Mims PN, Roman RJ. Knockdown of Add3 impairs the myogenic response of renal afferent arterioles and middle cerebral arteries. Am J Physiol Renal Physiol. 2017;312:F971–F981. doi: 10.1152/ajprenal.00529.2016

32. Gong CX, Dai CL, Liu F, Iqbal K. Multi-Targets: An Unconventional Drug Development Strategy for Alzheimer’s Disease. Front Aging Neurosci. 2022;14:837649. doi: 10.3389/fnagi.2022.837649

33. Duggan MR, Morgan DG, Price BR, Rajbanshi B, Martin-Peña A, Tansey MG, Walker KA. Immune modulation to treat Alzheimer’s disease. Mol Neurodegener. 2025;20:39. doi: 10.1186/s13024-025-00828-x

34. Yamashita T, Abe K. Update on Antioxidant Therapy with Edaravone: Expanding Applications in Neurodegenerative Diseases. Int J Mol Sci. 2024;25. doi: 10.3390/ijms25052945

35. Lomelí Martínez SM, Pacheco Moisés FP, Bitzer-Quintero OK, Ramírez-Jirano J, Delgado-Lara DLC, Cortés Trujillo I, Torres Jasso JH, Salazar-Flores J, Torres-Sánchez ED. Effect of N-Acetyl Cysteine as an Adjuvant Treatment in Alzheimer’s Disease. Brain Sci. 2025;15. doi: 10.3390/brainsci15020164

36. Abdalla M, Ibrahim M, Alkorbi N, Alkuwari S, Pedersen S, Rathore HA. Epoxide Hydrolase Inhibitors for the Treatment of Alzheimer’s Disease and Other Neurological Disorders: A Comprehensive Review. Biomedicines. 2025;13. doi: 10.3390/biomedicines13092073

37. Gregory A, Tang C, Fan F. Soluble epoxide hydrolase: a next-generation drug target for Alzheimer’s disease and related dementias. Neural Regen Res. 2025;20:2585–2586. doi: 10.4103/nrr.Nrr-d-24-00503

38. Fang X, Border JJ, Zhang H, Challagundla L, Kaur J, Hwang SH, Hammock BD, Fan F, Roman RJ. A Soluble Epoxide Hydrolase Inhibitor Improves Cerebrovascular Dysfunction, Neuroinflammation, Amyloid Burden, and Cognitive Impairments in the hAPP/PS1 TgF344-AD Rat Model of Alzheimer’s Disease. Int J Mol Sci. 2025;26. doi: 10.3390/ijms26062433

39. Morgan GC, Gregory A, Tang C, Hwang SH, Border JJ, Xu J, Liu Y, Bai S, Lee TJ, Cantwell C, et al. PTUPB improves cognitive function in Alzheimer’s disease associated with enhancing cerebral vascular myogenic response and attenuating vascular remodeling. Geroscience. 2026. doi: 10.1007/s11357-026-02140-y

40. Rosu GC, Catalin B, Balseanu TA, Laurentiu M, Claudiu M, Kumar-Singh S, Daniel P. Inhibition of Aquaporin 4 Decreases Amyloid Aβ40 Drainage Around Cerebral Vessels. Mol Neurobiol. 2020;57:4720–4734. doi: 10.1007/s12035-020-02044-8

41. Huber VJ, Igarashi H, Ueki S, Kwee IL, Nakada T. Aquaporin-4 facilitator TGN-073 promotes interstitial fluid circulation within the blood-brain barrier: [17O]H2O JJVCPE MRI study. Neuroreport. 2018;29:697–703. doi: 10.1097/wnr.0000000000000990

42. Tang C, Zhang H, Border JJ, Liu Y, Fang X, Jefferson JR, Gregory A, Johnson C, Lee TJ, Bai S, et al. Impact of knockout of dual-specificity protein phosphatase 5 on structural and mechanical properties of rat middle cerebral arteries: implications for vascular aging. Geroscience. 2024;46:3135–3147. doi: 10.1007/s11357-024-01061-y

43. Gregory AP, Morgan GC, Tang C, Zhang H, Border JJ, Fang X, Roman RJ, Fan F. SGLT2 Inhibition Protects Cognitive Function in Alzheimer’s Disease by Enhancing Brain Perfusion, Independent of Glucose Control. In: Alzheimers Dement. Alzheimers Dement: Alzheimers Dement. 2025 Dec 23;21(Suppl 1):e096815. doi: 10.1002/alz70855_096815. eCollection 2025 Dec.; 2026.

44. Wang S, Jiao F, Border JJ, Fang X, Crumpler RF, Liu Y, Zhang H, Jefferson J, Guo Y, Elliott PS, et al. Luseogliflozin, a sodium-glucose cotransporter-2 inhibitor, reverses cerebrovascular dysfunction and cognitive impairments in 18-mo-old diabetic animals. Am J Physiol Heart Circ Physiol. 2022;322:H246–H259. doi: 10.1152/ajpheart.00438.2021

45. Pardeshi R, Bolshette N, Gadhave K, Arfeen M, Ahmed S, Jamwal R, Hammock BD, Lahkar M, Goswami SK. Docosahexaenoic Acid Increases the Potency of Soluble Epoxide Hydrolase Inhibitor in Alleviating Streptozotocin-Induced Alzheimer’s Disease-Like Complications of Diabetes. Front Pharmacol. 2019;10:288. doi: 10.3389/fphar.2019.00288

46. Wu J, Zhao Y, Fan Z, Chen Q, Chen J, Sun Y, Jiang X, Xiao Q. Soluble epoxide hydrolase inhibitor protects against blood-brain barrier dysfunction in a mouse model of type 2 diabetes via the AMPK/HO-1 pathway. Biochem Biophys Res Commun. 2020;524:354–359. doi: 10.1016/j.bbrc.2020.01.085

47. Fan F, Muroya Y, Roman RJ. Cytochrome P450 eicosanoids in hypertension and renal disease. Curr Opin Nephrol Hypertens. 2015;24:37–46. doi: 10.1097/MNH.0000000000000088

48. Imig JD. Epoxyeicosatrienoic Acids and 20-Hydroxyeicosatetraenoic Acid on Endothelial and Vascular Function. Adv Pharmacol. 2016;77:105–141. doi: 10.1016/bs.apha.2016.04.003

49. Sura P, Sura R, Enayetallah AE, Grant DF. Distribution and expression of soluble epoxide hydrolase in human brain. J Histochem Cytochem. 2008;56:551–559. doi: 10.1369/jhc.2008.950659

50. Domingues MF, Callai-Silva N, Piovesan AR, Carlini CR. Soluble Epoxide Hydrolase and Brain Cholesterol Metabolism. Front Mol Neurosci. 2019;12:325. doi: 10.3389/fnmol.2019.00325

51. Oguro A, Fujita N, Imaoka S. Regulation of soluble epoxide hydrolase (sEH) in mice with diabetes: high glucose suppresses sEH expression. Drug Metab Pharmacokinet. 2009;24:438–445. doi: 10.2133/dmpk.24.438

52. Minaz N, Razdan R, Hammock BD, Goswami SK. An inhibitor of soluble epoxide hydrolase ameliorates diabetes-induced learning and memory impairment in rats. Prostaglandins Other Lipid Mediat. 2018;136:84–89. doi: 10.1016/j.prostaglandins.2018.05.004

53. Wu J, Fan Z, Zhao Y, Chen Q, Xiao Q. Inhibition of soluble epoxide hydrolase (sEH) protects hippocampal neurons and reduces cognitive decline in type 2 diabetic mice. Eur J Neurosci. 2021;53:2532–2540. doi: 10.1111/ejn.15150

54. Rose TE, Morisseau C, Liu JY, Inceoglu B, Jones PD, Sanborn JR, Hammock BD. 1-Aryl-3-(1-acylpiperidin-4-yl)urea inhibitors of human and murine soluble epoxide hydrolase: structure-activity relationships, pharmacokinetics, and reduction of inflammatory pain. J Med Chem. 2010;53:7067–7075. doi: 10.1021/jm100691c

55. Ostermann AI, Herbers J, Willenberg I, Chen R, Hwang SH, Greite R, Morisseau C, Gueler F, Hammock BD, Schebb NH. Oral treatment of rodents with soluble epoxide hydrolase inhibitor 1-(1-propanoylpiperidin-4-yl)-3-[4-(trifluoromethoxy)phenyl]urea (TPPU): Resulting drug levels and modulation of oxylipin pattern. Prostaglandins Other Lipid Mediat. 2015;121:131–137. doi: 10.1016/j.prostaglandins.2015.06.005

56. Pabbidi MR, Mazur O, Fan F, Farley JM, Gebremedhinm D, Harder DR, Roman RJ. Enhanced large conductance K+ channel (BK) activity contributes to the impaired myogenic response in the cerebral vasculature of Fawn Hooded Hypertensive rats. Am J Physiol Heart Circ Physiol. 2014;2014 Apr 1;306(7):H989-H1000.

57. Burke M, Pabbidi M, Fan F, Ge Y, Liu R, Williams JM, Sarkis A, Lazar J, Jacob HJ, Roman RJ. Genetic basis of the impaired renal myogenic response in FHH rats. Am J Physiol Renal Physiol. 2013;304:F565–577. doi: 10.1152/ajprenal.00404.2012

58. Genitsaridi I, Salpea P, Salim A, Sajjadi SF, Tomic D, James S, Thirunavukkarasu S, Issaka A, Chen L, Basit A, et al. 11th edition of the IDF Diabetes Atlas: global, regional, and national diabetes prevalence estimates for 2024 and projections for 2050. Lancet Diabetes Endocrinol. 2026;14:149–156. doi: 10.1016/s2213-8587(25)00299-2

59. Mansour GK, Bolgova O, Hajjar AW, Mavrych V. Neurovascular Dysfunction and Glymphatic Impairment: An Unexplored Therapeutic Frontier in Neurodegeneration. Int J Mol Sci. 2025;26. doi: 10.3390/ijms262411843

60. Wolters FJ, Zonneveld HI, Hofman A, van der Lugt A, Koudstaal PJ, Vernooij MW, Ikram MA. Cerebral Perfusion and the Risk of Dementia: A Population-Based Study. Circulation. 2017;136:719–728. doi: 10.1161/circulationaha.117.027448

61. Fan F, Booz GW, Roman RJ. Aging diabetes, deconstructing the cerebrovascular wall. Aging (Albany NY). 2021;13:9158–9159. doi: 10.18632/aging.202963

62. Fang X, Zhang J, Roman RJ, Fan F. From 1901 to 2022, how far are we from truly understanding the pathogenesis of age-related dementia? GeroScience. 2022;44:1879–1883. doi: 10.1007/s11357-022-00591-7

63. Lee KSS, Yang J, Niu J, Ng CJ, Wagner KM, Dong H, Kodani SD, Wan D, Morisseau C, Hammock BD. Drug-Target Residence Time Affects in Vivo Target Occupancy through Multiple Pathways. ACS Cent Sci. 2019;5:1614–1624. doi: 10.1021/acscentsci.9b00770

64. Ghosh A, Comerota MM, Wan D, Chen F, Propson NE, Hwang SH, Hammock BD, Zheng H. An epoxide hydrolase inhibitor reduces neuroinflammation in a mouse model of Alzheimer’s disease. Sci Transl Med. 2020;12. doi: 10.1126/scitranslmed.abb1206

65. Ulu A, Inceoglu B, Yang J, Singh V, Vito S, Wulff H, Hammock BD. Inhibition of soluble epoxide hydrolase as a novel approach to high dose diazepam induced hypotension. J Clin Toxicol. 2016;6. doi: 10.4172/2161-0495.1000300

66. Moshal KS, Zeldin DC, Sithu SD, Sen U, Tyagi N, Kumar M, Hughes WM, Jr., Metreveli N, Rosenberger DS, Singh M, et al. Cytochrome P450 (CYP) 2J2 gene transfection attenuates MMP-9 via inhibition of NF-kappabeta in hyperhomocysteinemia. J Cell Physiol. 2008;215:771–781. doi: 10.1002/jcp.21356

67. Ambrosini E, Aloisi F. Chemokines and glial cells: a complex network in the central nervous system. Neurochem Res. 2004;29:1017–1038. doi: 10.1023/b:nere.0000021246.96864.89

68. Harrison IF, Ismail O, Machhada A, Colgan N, Ohene Y, Nahavandi P, Ahmed Z, Fisher A, Meftah S, Murray TK, et al. Impaired glymphatic function and clearance of tau in an Alzheimer’s disease model. Brain. 2020;143:2576–2593. doi: 10.1093/brain/awaa179

69. Attwell D, Buchan AM, Charpak S, Lauritzen M, Macvicar BA, Newman EA. Glial and neuronal control of brain blood flow. Nature. 2010;468:232–243. doi: 10.1038/nature09613

70. Bhat NR, Fan F. Adenovirus infection induces microglial activation: involvement of mitogen-activated protein kinase pathways. Brain Res. 2002;948:93–101. doi: 10.1016/s0006-8993(02)02953-0

